# *Arabidopsis* PFA-DSP-type phosphohydrolases target specific inositol pyrophosphate messengers

**DOI:** 10.1101/2022.03.01.482514

**Authors:** Philipp Gaugler, Robin Schneider, Guizhen Liu, Danye Qiu, Jonathan Weber, Jochen Schmid, Nikolaus Jork, Markus Häner, Kevin Ritter, Nicolás Fernández-Rebollo, Ricardo F.H. Giehl, Minh Nguyen Trung, Ranjana Yadav, Dorothea Fiedler, Verena Gaugler, Henning J. Jessen, Gabriel Schaaf, Debabrata Laha

## Abstract

Inositol pyrophosphates are signaling molecules containing at least one phosphoanhydride bond that regulate a wide range of cellular processes in eukaryotes. With a cyclic array of phosphate esters and diphosphate groups around *myo*-inositol, these molecular messengers possess the highest charge density found in nature. Recent work deciphering inositol pyrophosphate biosynthesis in *Arabidopsis* revealed important functions of these messengers in nutrient sensing, hormone signaling and plant immunity. However, despite the rapid hydrolysis of these molecules in plant extracts, very little is known about the molecular identity of the phosphohydrolases that convert these messengers back to their inositol polyphosphate precursors. Here, we investigate whether *Arabidopsis* Plant and Fungi Atypical Dual Specificity Phosphatases (PFA-DSP1-5) catalyze inositol pyrophosphate phosphohydrolase activity. We find that recombinant proteins of all five *Arabidopsis* PFA-DSP homologs display phosphohydrolase activity with a high specificity for the 5-β-phosphate of inositol pyrophosphates. We further show that heterologous expression of *Arabidopsis* PFA-DSP1-5 rescues wortmannin-sensitivity and deranged inositol pyrophosphate homeostasis caused by the deficiency of the PFA-DSP-type inositol pyrophosphate phosphohydrolase Siw14 in yeast. Heterologous expression in *Nicotiana benthamiana* leaves provided evidence that *Arabidopsis* PFA-DSP1 also displays 5-β-phosphate specific inositol pyrophosphate phosphohydrolase activity *in planta*. Our findings lay the biochemical basis and provide the genetic tools to uncover the roles of inositol pyrophosphates in plant physiology and plant development.

## INTRODUCTION

Inositol pyrophosphates (PP-InsPs), such as InsP_7_ and InsP_8_, are molecules derived from *myo*-inositol (Ins) esterified with unique patterns of monophosphates (P) and diphosphates (PP), and have been described as versatile messengers in yeast, amoeba, and animal cells ^*1-4*^. With recent discoveries that PP-InsPs regulate nutrient sensing and immunity in plants, these molecules are a novel focus of research in plant physiology ^*5-13*^. The synthesis of PP-InsPs is partially conserved in eukaryotes, with some important distinctions in plants. In baker’s yeast and mammals, 5-InsP_7_ is synthesized by Kcs1/IP6K-type proteins whereas Vip1/PPIP5K-type kinases phosphorylate the C1 position of both InsP_6_ (also termed phytic acid) and 5-InsP_7_, generating 1-InsP_7_ and 1,5-InsP_8_, respectively ^*14-17*^. In plants, detection, quantification and characterization of PP-InsPs have been challenging due to the low abundance of these molecules and their susceptibility to hydrolytic activities during extraction ^*18, 19*^. Employing [^3^H] *myo*-inositol labelling and subsequent analysis of plant extracts by strong-anion exchange high performance liquid chromatography (SAX-HPLC) allowed the detection of PP-InsPs in different plant species ^*5, 20-22*^. The recent development of capillary electrophoresis (CE) electrospray ionization mass spectrometry (ESI-MS), has enabled the detection and quantification of many InsP and PP-InsP isomers in various cell extracts including all InsP_7_ isomers, except enantiomers (labeled e.g. as 1/3 or 4/6 InsP_7_) ^*23*^. Similar to yeast and mammals, the *Arabidopsis* PPIP5K isoforms, VIH1 and VIH2 catalyse the synthesis of InsP_8_ ^*5, 10*^ and are likely involved in the synthesis of 1/3-InsP_7_ ^*11*^. However, Kcs1/IP6K-type proteins are absent in land plants. The question of how plants synthesize 5-InsP_7_ has been partially solved by work on *Arabidopsis* inositol (1,3,4) triphosphate 5/6 kinases ITPK1 and ITPK2. Notably, ITPK1 and ITPK2 were reported to catalyse the synthesis of 5-InsP_7_ from InsP_6_ *in vitro* ^*24-26*^ and consequently *itpk1* mutant plants display reduced 5-InsP_7_ levels ^*11, 13*^.

In *Arabidopsis*, disturbances in the synthesis of InsP_7_ and/or InsP_8_ result in defective signaling of the plant hormones jasmonate ^*5, 6*^ and auxin ^*13*^, as well as defects in salicylic acid-dependent plant immunity ^*12*^ and impaired phosphate (P_i_) homeostasis ^*9-11, 27*^. In the case of auxin and jasmonate perception, 5-InsP_7_ and InsP_8_, respectively, are proposed to function as co-ligands of the respective receptor complexes ^*5, 6, 13*^. The role of PP-InsPs in P_i_ signaling are related to their ability to bind to SPX proteins, which act as receptors for these messenger molecules ^*7, 9, 10, 28-30*^. InsP_8_ has been found as the preferred ligand for stand-alone SPX proteins *in vivo* ^*9, 10, 28*^. InsP_8_-bound SPX receptors inactivate the MYB-type transcription factors PHR1 and PHL1, which control the expression of a majority of P_i_ starvation-induced (PSI) genes to regulate various metabolic and developmental adaptations induced by P_i_ deficiency ^*28, 31-33*^. The tissue levels of various PP-InsPs, including 5-InsP_7_ and InsP_8_, respond sensitively to the plant’s P_i_ status ^*9, 11*^, suggesting that their synthesis and degradation are tightly regulated. While the steps involved in the synthesis of PP-InsPs in plants are now better understood, still little is known about how these molecules are degraded.

Vip1/PPIP5Ks are bifunctional enzymes that harbour an N-terminal ATP-grasp kinase domain and a C-terminal phosphatase domain conserved in yeast, animals and plants ^*5, 10, 15, 17, 34*^. *In vitro*, the phosphatase domain of *Arabidopsis* PPIP5K VIH2 hydrolyses PP-InsPs to InsP_6_ ^*10*^, similar to the respective C-terminal domains of fission yeast and mammalian PPIP5Ks ^*35, 36*^. Although *Arabidopsis* ITPK1 harbors no phosphatase domain, under conditions of low adenylate charge it can shift its activity *in vitro* from kinase to ADP phosphotransferase activity using 5-InsP_7_ but no other InsP_7_ isomer ^*11, 26*^. Apart from relying on the reversible activities of ITPK1 and Vip1/PPIP5Ks, the degradation of PP-InsPs may also be controlled by specialized phosphohydrolases.

In mammalian cells, diphosphoinositol polyphosphate phosphohydrolases (DIPPs), members of the nudix hydrolase family, have been shown to catalyse hydrolysis of the diphosphate groups of InsP_7_ and InsP_8_ at the C1 and C5 position ^*4, 37, 38*^. The baker’s yeast genome encodes a single homologue of mammalian DIPP1, named diadenosine and diphosphoinositol polyphosphate phosphohydrolase (DDP1), which hydrolyses various substrates including diadenosine polyphosphates, 5-InsP_7_ and InsP_8_, but has a preference for inorganic polyphosphates (poly-P) and for the β-phosphate of 1-InsP_7_ ^*39-41*^. In addition, baker’s yeast has an unrelated PP-InsP phosphohydrolase, Siw14 (also named Oca3) with a high specificity for the β-phosphate at position C5 of 5-InsP_7_ ^*42, 43*^. This enzyme is a member of the Plant and Fungi Atypical Dual Specificity Phosphatases (PFA-DSPs) that belong to a large family of protein tyrosine phosphatases (PTPs) ^*43-45*^.

Blast search analyses revealed that the *Arabidopsis thaliana* genome encodes five PFA-DSPs, with AtPFA-DSP1 sharing 61% identity and 76% similarity with yeast Siw14 ^*44, 45*^. X-ray crystallography revealed that the protein adopts an α/β fold typical for cysteine phosphatases, with the predicted catalytic cysteine (Cys150) residing at the bottom of a positively-charged pocket ^*44, 46*^. Of a number of putative phosphatase substrates tested, recombinant AtPFA-DSP1 displayed the highest activity against inorganic polyphosphate, as well as against deoxyribo- and ribonucleoside triphosphates, and less activity against phosphotyrosine-containing peptides and phosphoinositides ^*46*^. Here, we investigated whether *Arabidopsis* PFA-DSPs might function as PP-InsP phosphohydrolases.

## Results and Discussion

### *Arabidopsis* PFA-DSP proteins display *in vitro* PP-InsP phosphohydrolase activity with high specificity for 5-InsP_7_

To explore a potential role of *Arabidopsis* PFA-DSP proteins in PP-InsP hydrolysis, we firstly generated translational fusions of PFA-DSPs with an N-terminal hexahistidine tag followed by a maltose binding protein (MBP) and expressed recombinant proteins in bacteria. Corresponding His-MBP-Siw14 and free His-MBP constructs were generated as controls. All constructs allowed the purification of soluble recombinant proteins (Figure S1). We then tested potential PP-InsP phosphohydrolase activities of PFA-DSP1 with 1-InsP_7_ or 5-InsP_7_ in the presence of various divalent cations. Notably, PFA-DSP1 failed to catalyze the hydrolysis of 1-InsP_7_ or 5-InsP_7_ in the presence of Mn^2+^, Ca^2+^ or Zn^2+^. However, in the presence of the cytoplasmic prevalent cation Mg^2+^, PFA-DSP1 displayed a robust hydrolytic activity against 5-InsP_7_, likely resulting in the generation of InsP_6_, as deduced from the mobility of the reaction product separated by polyacrylamide gel electrophoresis (PAGE) and visualized by toluidine blue staining (Figure 1A). CE-ESI-MS analysis of the reaction product spiked with a [^13^C_6_] InsP_6_ standard confirmed that the resulting product indeed had the migration behavior and the mass of phytic acid (Figure 1B). In contrast, 1-InsP_7_ was largely resistant to PFA-DSP1 also in the presence of Mg^2+^ (Figure 1A). In the absence of divalent cations (i.e. in buffer not supplemented with divalent cations but instead supplemented with EDTA, a condition unlikely to represent any cellular condition) both InsP_7_ isomers were hydrolyzed to InsP_6_, as deduced from the mobility of the reaction product by PAGE (Figure 1A).

**Figure 1:**
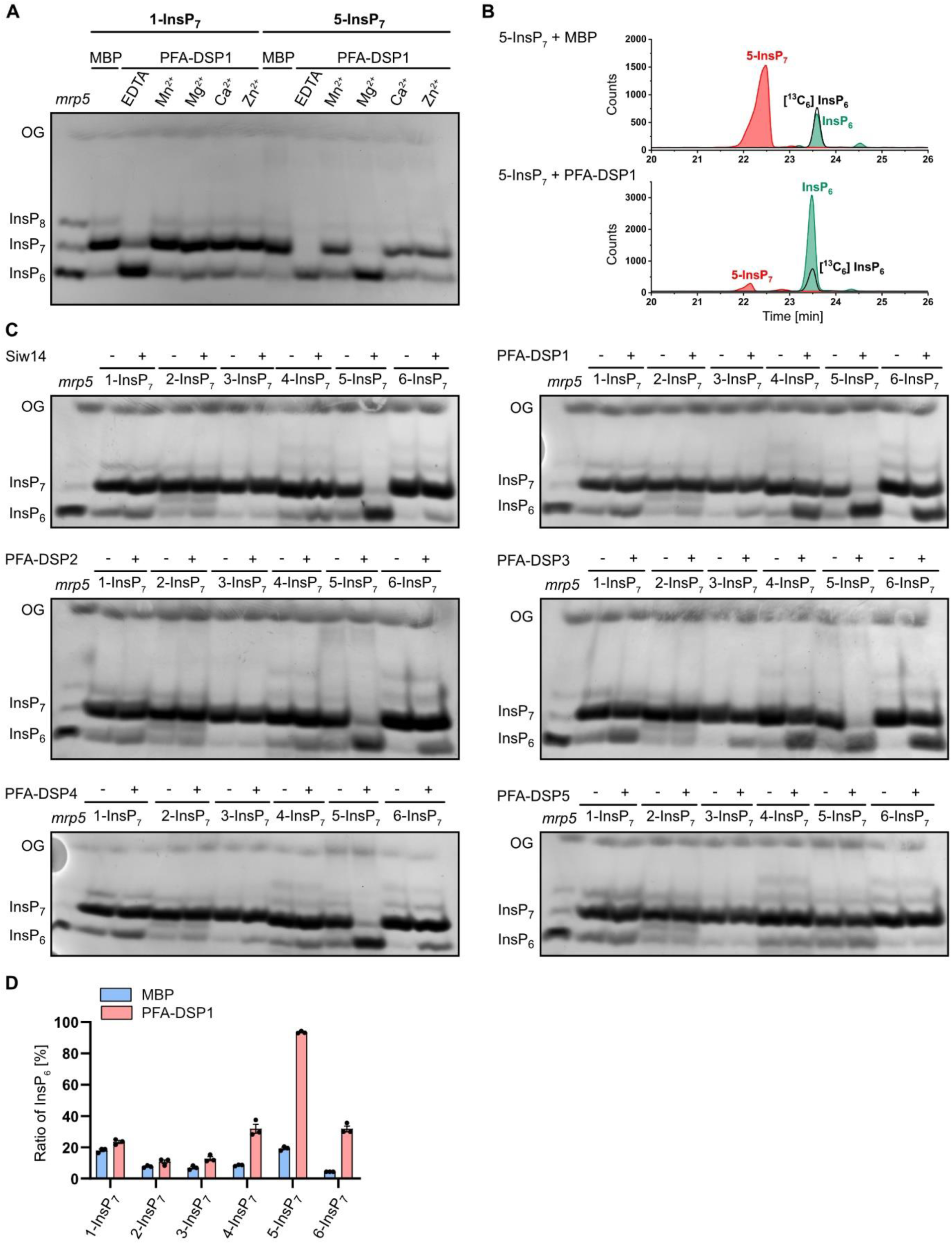
*In vitro, Arabidopsis* PFA-DSPs display Mg^2+^-dependent PP-InsP phosphohydrolase activity with high specificity for 5-InsP_7_. Recombinant His-MBP-DSPs and His-MBP-Siw14 were incubated with 0.33 mM InsP_7_ for 1 h at 22°C. His-MBP served as a negative control. (A) 0.4 μM His-MBP-DSP1 was incubated with 1-InsP_7_ or 5-InsP_7_ and either 1 mM EDTA, MnCl_2_, MgCl_2_, CaCl_2_ or ZnCl_2_ as indicated. The reaction products were then separated by 33 % PAGE and visualized by toluidine blue. (B-D) The InsP_7_ phosphohydrolase activity of approximately 0.4 μM His-MBP-DSPs and His-MBP-Siw14 was analyzed in presence of 1 mM MgCl_2_. The reaction products were then (B, D) spiked with isotopic standards mixture ([^13^C_6_]1,5-InsP_8_, [^13^C_6_]5-InsP_7_, [^13^C_6_]1-InsP_7_, [^13^C_6_] InsP_6_, [^13^C_6_]2-OH InsP_5_) and subjected to CE-ESI-MS analyses or (C) separated by 33 % PAGE and visualized by toluidine blue/DAPI staining. (D) Data represent means ± SEM (n = 3). Representative extracted-ion electropherograms are shown in Figure S2.

We then tested the hydrolytic activities of the *Arabidopsis* PFA-DSP homologs with all six ‘simple’, synthetic InsP_7_ isomers and with the two enantiomeric InsP_8_ isomers 1,5-InsP_8_ and 3,5-InsP_8_ in the presence of Mg^2+^. Of note, *myo*-inositol is a meso compound with a mirror plane dissecting the C2 and C5 position. Derivatives differentially (pyro)phosphorylated at the C1 and C3 position, as well as at the C4 and C6 position are enantiomeric forms that can only be distinguished in the presence of appropriate chiral selectors ^*47, 48*^. Yeast Siw14 and all *Arabidopsis* PFA-DSPs with the exception of PFA-DSP5, displayed robust activity with a high specificity towards 5-InsP_7_ (Figure 1C), confirming earlier reports that 5-InsP_7_ is a preferred substrate for yeast Siw14 as compared to 1-InsP_7_ ^*42, 43*^. PFA-DSP1-4 and Siw14 also displayed partial hydrolytic activities against the enantiomers 4-InsP_7_ and 6-InsP_7_, as well as very weak hydrolytic activities against enantiomeric 1-InsP_7_ and 3-InsP_7_ (Figure 1C). The latter activities were more pronounced in PFA-DSP1 and PFA-DSP3 as compared to Siw14 and PFA-DSP2. As for 5-InsP_7_, the reaction products with the other InsP_7_ isomers had the mass and the migration behavior of the InsP_6_ isomer phytic acid, as deduced from CE-ESI-MS analyses (Figures 1D and S2).

Notably, PFA-DSP5 only showed very weak activities at the 0.4 μM concentration tested in our assay. However, when the reaction time was extended from 1 h to 2 h and the enzyme concentration was increased to 2 μM, PFA-DSP5 displayed robust activity with a substrate specificity similar to PFA-DSP1-4 and yeast Siw14, with a high selectivity for 5-InsP_7_ and only weak hydrolytic activities against 4-InsP_7_ and 6-InsP_7_ (Figure 2A). Notably, the meso InsP_7_ isomer 2-InsP_7_ was completely resistant to Siw14 or any of the *Arabidopsis* PFA-DSP proteins under the assay conditions. This was also the case in the absence of divalent cations (i.e. in buffer not supplemented with divalent cations but instead supplemented with EDTA), where Siw14 and *Arabidopsis* PFA-DSP1-4 failed to hydrolyze 2-InsP_7_ to a significant extent while all other InsP_7_ isomers were at least partially converted to an InsP isomer with the mobility of phytic acid (Figures S3 and S4). Even after a 24 h-long incubation with *Arabidopsis* PFA-DSP1, 2-InsP_7_ remained largely resistant to hydrolysis. In contrast, all other PP-InsP_7_ isomers were hydrolyzed to InsP_6_ under these conditions, as revealed by PAGE and CE-ESI-MS analyses (Figures 2B,C and S5). Corresponding control reactions that were supplemented with 5-InsP_7_ after 23 h validated the activity of *Arabidopsis* PFA-DSP1 after such long incubation times (Figures 2C and S6). These spiking experiments also rules out the possibility that 2-InsP_7_ contained a contaminant that inhibits PFA-DSP-dependent hydrolysis, as 5-InsP_7_ was still efficiently hydrolyzed in the presence of 2-InsP_7_.

**Figure 2:**
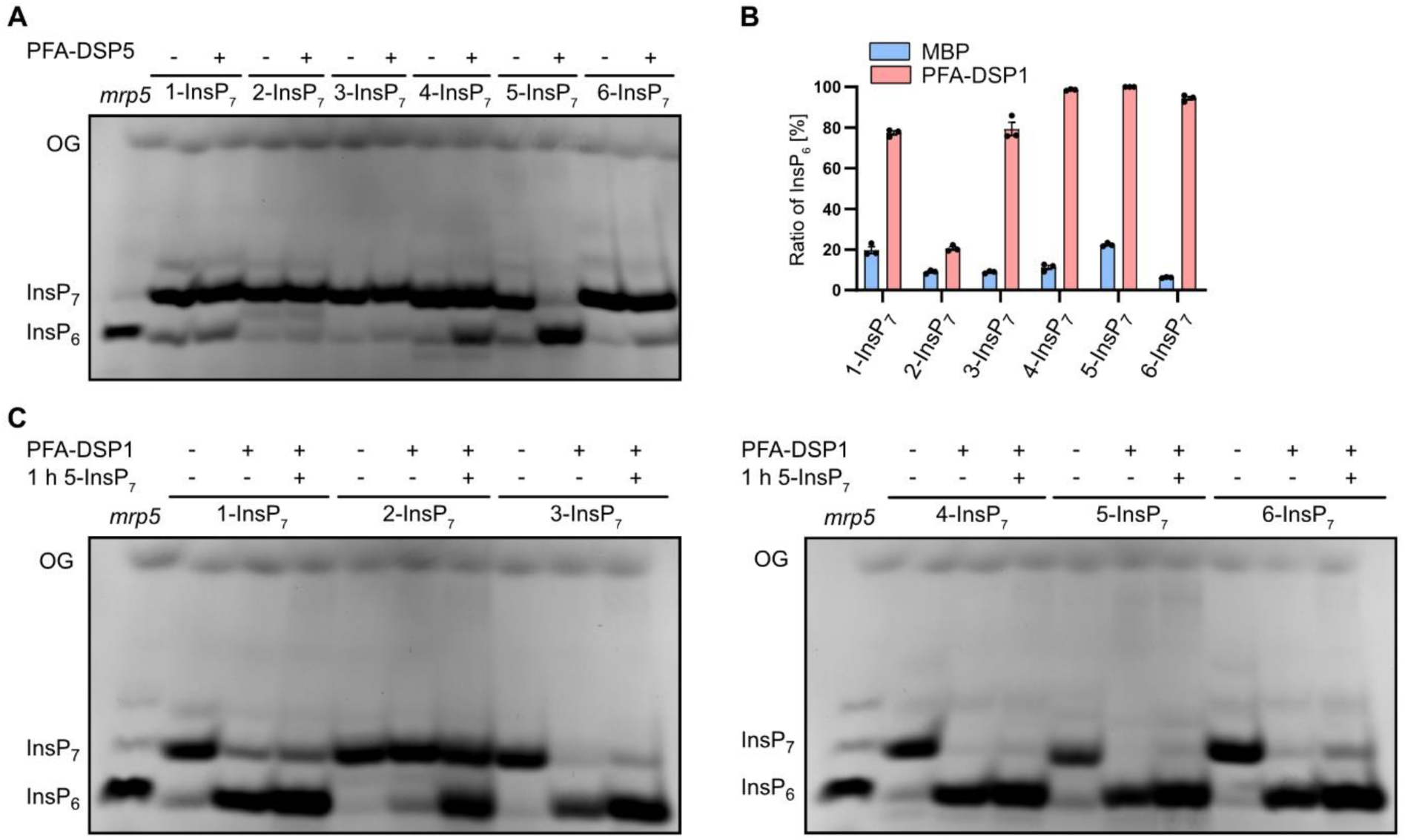
Under prolonged incubation times and/or at increased protein concentrations, selected *Arabidopsis* PFA-DSPs hydrolyze various InsP_7_ isomers *in vitro*, except for 2-InsP_7_. Recombinant His-MBP-DSPs or His-MBP-Siw14 were incubated with 1 mM MgCl_2_ at 22°C. His-MBP served as a negative control. (A) 2 μM recombinant His-MBP-DSP5 was incubated with 0.33 mM InsP_7_ isomers for 2 h and the reaction product was separated by 33 % PAGE and visualized with toluidine blue. (B, C) 0.4 μM PFA-DSP1 was incubated with 0.33 mM InsP_7_ for 24 h. To ensure that PFA-DSP1 is active during the whole incubation time, after 23 h, 0.33 mM 5-InsP_7_ was added to a replicate and incubated for another 1 h. (B) An aliquot of the reaction product was spiked with an isotopic standards mixture ([^13^C_6_]1,5-InsP_8_, [^13^C_6_]5-InsP_7_, [^13^C_6_]1-InsP_7_, [^13^C_6_] InsP_6_, [^13^C_6_]2-OH InsP_5_) and subjected to CE-ESI-MS analyses. Data represent means ± SEM (n = 3). (C) Another reaction was separated by 33 % PAGE and visualized with toluidine blue. Representative extracted-ion electropherograms are shown in Figure S5.

Finally, we tested whether the enantiomeric InsP_8_ isomers 1,5-InsP_8_ and 3,5-InsP_8_ serve as substrates for PFA-DSPs. As reported earlier for Siw14 ^*43*^, PFA-DSP1-4 hydrolyzed 1,5-InsP_8_ to an InsP_7_ isomer based on the mobility of the reaction product in PAGE analyses (Figure 3A). Also the enantiomeric 3,5-InsP_8_ was efficiently hydrolyzed by Siw14 and PFA-DSP1-4 (Figure 3A) and CE-ESI-MS analysis of the reaction products showed the migration behavior and the mass of 1/3-InsP_7_ (Figures 3B,C and S7).

**Figure 3:**
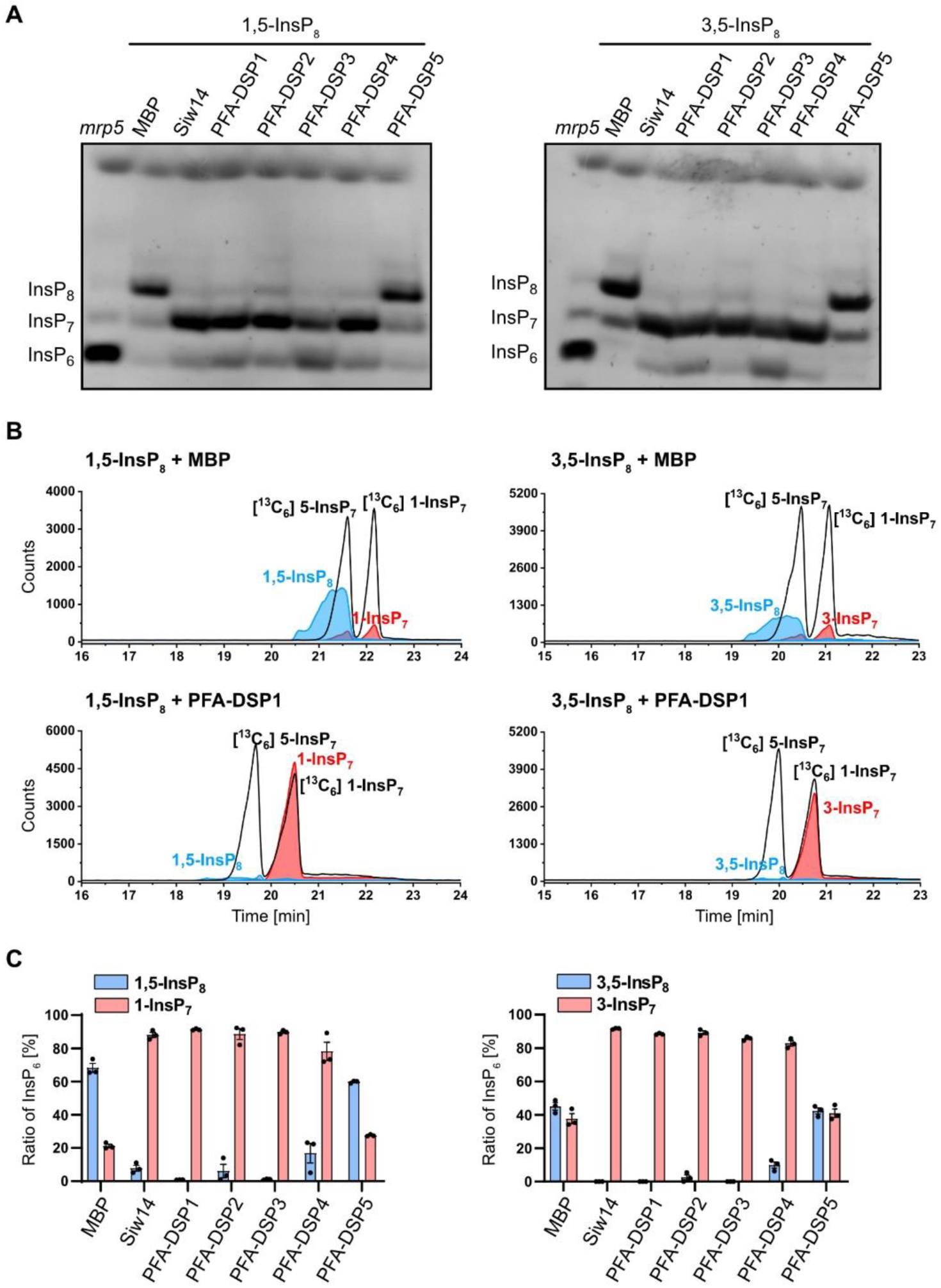
*In vitro, Arabidopsis* PFA-DSPs display robust 1/3,5-InsP_8_ phosphohydrolase activity. (A) Approximately 0.4 μM His-MBP-DSPs and His-MBP-Siw14 were incubated with 0.33 mM 1,5-InsP_8_ or 3,5-InsP_8_ for 1 h, in presence of 1 mM MgCl_2_, and analyzed by PAGE and subsequent toluidine blue staining. (B, C) A second and third reaction were spiked with isotopic standards mixture ([^13^C_6_]1,5-InsP_8_, [^13^C_6_]5-InsP_7_, [^13^C_6_]1-InsP_7_, [^13^C_6_] InsP_6_, [^13^C_6_]2-OH InsP_5_) and subjected to CE-ESI-MS analyses. (C) Data represent means ± SEM (n = 3). Representative extracted-ion electropherograms are shown in Figure S7.

Altogether, these findings reveal that *Arabidopsis* PFA-DSP proteins, as well as Siw14 have a high specificity for the 5-β-phosphate of 5-InsP_7_, 1,5-InsP_8_ and 3,5-InsP_8_, and a weak activity against the β-phosphates of 4-InsP_7_ and 6-InsP_7_, respectively. In contrast, InsP_6_ and 2-InsP_7_ are resistant to PFA-DSP-catalyzed hydrolysis.

### Heterologous expression of *Arabidopsis PFA-DSP* homologs complement yeast *siw14Δ*defects

To investigate the physiological consequences of *Arabidopsis* PFA-DSP activities *in vivo*, we carried out heterologous expression experiments in baker’s yeast. To this end, we first screened the *siw14Δ* yeast strain for phenotypes and identified a not yet described severe growth defect on media containing the fungal toxin wortmannin ^*49*^. Previous observations that *kcs1Δ* yeast cells are resistant to wortmannin ^*50*^ suggest that wortmannin sensitivity of *siw14Δ* yeast might be related to Kcs1-dependent PP-InsPs. The *siw14Δ* -associated growth defect was fully complemented by heterologous expression of either of the five *Arabidopsis PFA-DSP* homologs or of yeast *SIW14* from episomal plasmids under control of a *PMA1* promotor fragment (Figure 4A). To investigate the contribution of PFA-DSPs in InsP metabolism, we monitored InsP profiles using SAX-HPLC analyses of various [^3^H]-*myo*-inositol labelled yeast transformants. Of note, conventional SAX-HPLC analyses as employed here do not allow the discrimination of different InsP_7_ or InsP_8_ isomers ^*11, 48*^. Heterologous expression of PFA-DSPs in *siw14Δ* restored InsP_7_/InsP_6_ ratios to wild-type levels, indicating that *Arabidopsis* PFA-DSP proteins are functionally similar to Siw14 (Figure 4B).

**Figure 4:**
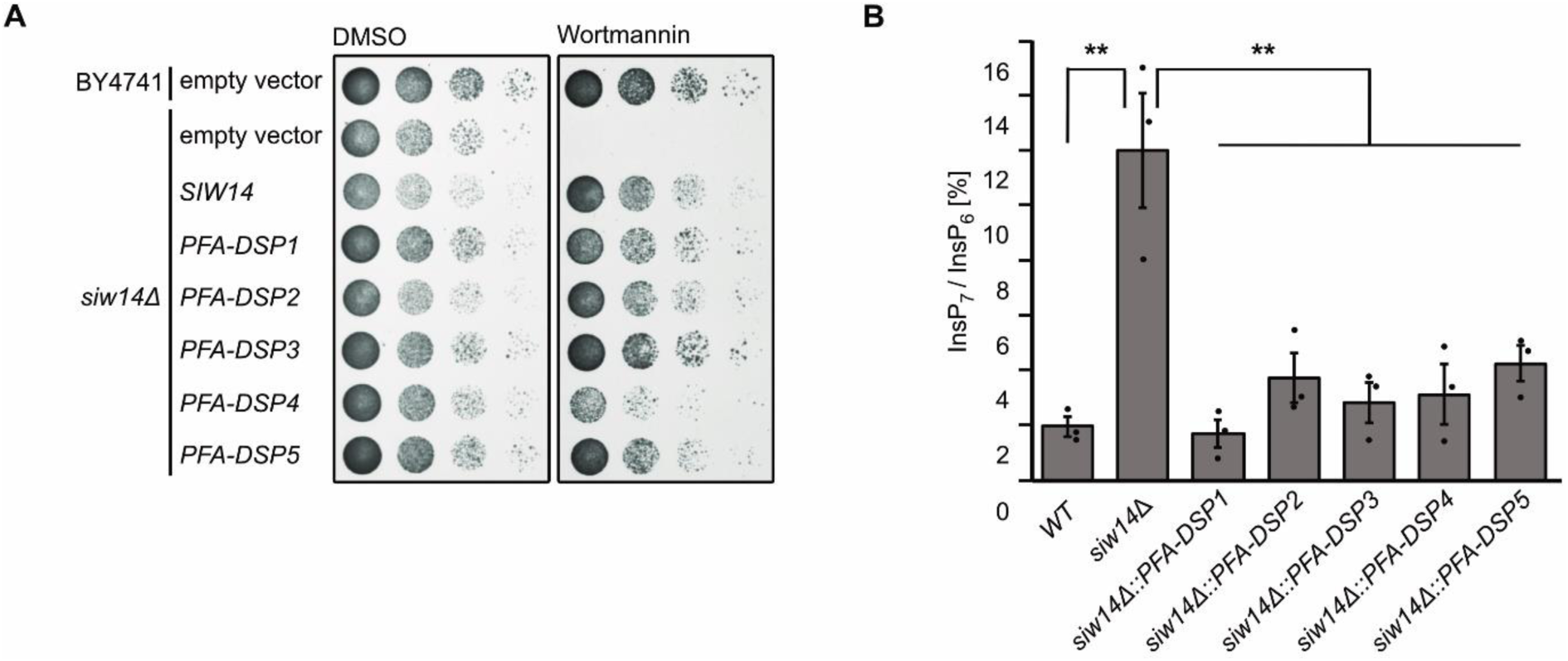
Heterologous expression of *Arabidopsis* PFA-DSPs complements *siw14Δ*-associated wortmannin sensitivity in yeast. (A) Growth complementation assays of an *siw14Δ* yeast strain. Wild-type yeast (BY4741) and an isogenic *siw14Δ* yeast mutant were transformed with either the empty episomal pDRf1-GW plasmid or with different pDRf1-GW plasmids carrying the respective *PFA-DSP* gene or *SIW14*. Yeast transformants were then spotted in 8-fold serial dilutions (starting from OD_600_ 1.0) onto selective media supplemented with either wortmannin or DMSO as control. Plates were incubated at 26 °C for 2 days before photographing. The yeast growth assay was repeated twice (n = 3) with similar results. (B) The relative amounts of InsP_7_ of wild-type yeast, *siw14Δ* and *siw14Δ* transformed with pDRf1-GW carrying the *DSP* genes are shown as InsP_7_/InsP_6_ ratios. InsP_6_ and InsP_7_ levels were determined by analysis of SAX-HPLC profiles using OriginPro 8. Data represent means ± SEM (n = 3). Asterisks indicate values that are significantly different from *siw14Δ* (according to Student’s *t* test, P<0.05 (*); P<0.01 (**); P<0.001 (***)).

Notably, the InsP_7_ signal was the only one consistently affected by the loss of *SIW14* and heterologous expression of any *PFA-DSP* gene (Figure S8). We generated variants of Siw14 or PFA-DSP1, in which the catalytic cysteine was replaced by a serine resulting in a C214S and a C150S substitution in Siw14 and PFA-DSP1, respectively, and observed that complementation of *siw14Δ*-associated growth defects of respective transformants requires the catalytic activity of these proteins (Figures S9A,B). The inability of catalytic dead mutants to complement *siw14Δ* -associated growth defects was not caused by compromised expression or protein stability of these variants, as confirmed by immunoblot analyses (Figure S9C). In agreement with the growth complementation assays, the catalytically inactive versions of Siw14 and PFA-DSP1 also failed to restore wild-type InsP_7_ levels in *siw14Δ* transformants (Figures S9D,E). These experiments suggest that *Arabidopsis* PFA-DSPs can substitute for endogenous Siw14 in yeast with respect to wortmannin tolerance and InsP_7_ homeostasis and that complementation of the *siw14Δ*-associated defects depends on the catalytic activity of these proteins.

### Growth defects of *siw14Δ* yeast on wortmannin require Kcs1-dependent 5-InsP_7_ synthesis

For a deeper understanding of the wortmannin phenotype of *siw14Δ* yeast, we investigated genetic interactions between Siw14 and different InsP kinases. We generated different double mutants with defects in Siw14 and the PP-InsP synthases Kcs1 and Vip1, and tested their performance on wortmannin-containing media (Figure 5A). Again, *siw14Δ* cells did not survive on media supplemented with 3 μM wortmannin, a defect that was fully complemented by the expression of *SIW14* under control of the endogenous promoter from a *CEN*-based single copy plasmid (Figure 5A). Growth of *vip1Δ* cells was comparable to wild-type yeast. In contrast, the *vip1Δ siw14Δ* double mutant showed a severe growth defect on media supplemented with wortmannin similar to single *siw14Δ* cells (Figure 5A). Like the *vip1Δ* yeast strain, a *kcs1Δ* strain did not show growth defects on media supplemented with wortmannin as compared to control media. In contrast, at increased concentrations we observed *kcs1Δ*-associated wortmannin resistance (Figure 5B), as reported earlier ^*50*^. Importantly, deletion of *KCS1* in *siw14Δ* cells rescued *siw14Δ*-associated wortmannin sensitivity since the resulting *kcs1Δ siw14Δ* double mutant yeast strain, despite growing overall weaker than the *kcs1Δ* single mutant strain, showed no increased sensitivity to wortmannin (Figure 5A). These findings indicate that the presence of Kcs1 is critical for the growth defects displayed by *siw14Δ* single mutant cells on wortmannin. We then investigated whether the presence of Kcs1 itself or of Kcs1-dependent PP-InsPs such as 5-InsP_7_ are relevant for *siw14Δ*-associated wortmannin sensitivity. To this end, we examined the phenotypes of *ipk2Δ* and of *ipk2Δ siw14Δ* yeast transformants. Both mutants lack IPK2, an inositol polyphosphate multikinase that sequentially phosphorylates Ins(1,4,5)P_3_ to Ins(1,3,4,5,6)P_5_ and is hence required for InsP_6_ and subsequent Kcs1-dependent 5-InsP_7_ or PP-InsP_4_ synthesis ^*50, 51*^. Neither of the strains showed growth defects on media supplemented with wortmannin as compared to the isogenic wild-type yeast strain, suggesting that also the loss of *IPK2* rescues *siw14Δ*-associated wortmannin sensitivity (Figure 5A). We further tested wortmannin sensitivity of *kcs1Δ* and *kcs1Δ siw14Δ* yeast transformants in a different genetic background and observed similar results (Figures 5B,C). Taken together, these results provide a causal link between Kcs1- (but not Vip1-) dependent PP-InsPs and *siw14Δ* -associated wortmannin sensitivity with Kcs1 and Siw14/PFA-DSPs playing antagonistic roles in regulating this sensitivity.

**Figure 5:**
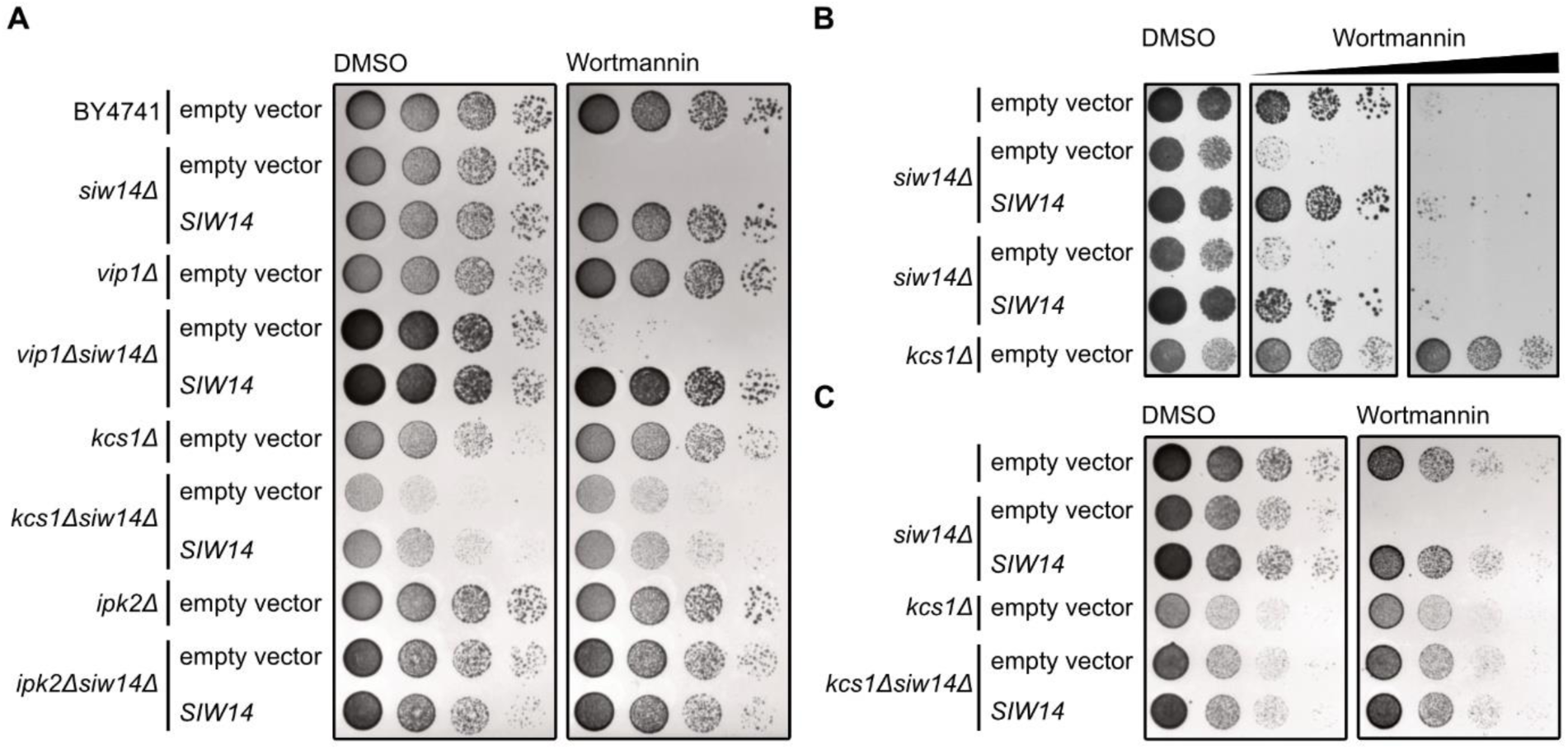
Yeast *siw14Δ*-associated wortmannin sensitivity requires Kcs1-dependent 5-InsP_7_. (A) A Wild-type yeast (BY4741), *siw14Δ, vip1Δ, kcs1Δ, ipk2Δ, vip1Δ siw14Δ, kcs1Δ siw14Δ* and *ipk2Δ siw14Δ* double mutant yeast strains were transformed with either empty YCplac33 vector or with a YCplac33 vector carrying a genomic fragment of *SIW14* including a 653 bp promoter and 5’UTR and a 289 bp terminator region. Yeast transformants were then spotted in 8-fold serial dilutions (starting from OD_600_ 1.0) onto selective media supplemented with either wortmannin or DMSO as control. Plates were incubated at 26 °C for 2 days before photographing. (B,C) Wild-type yeast (DDY1810), *siw14Δ, kcs1Δ* and (C) *kcs1Δ siw14Δ* double mutant yeast strains were transformed with either empty YCplac33 vector or with a YCplac33 vector carrying the genomic fragment of *SIW14*. The growth assay was performed as described for (A). All yeast growth assays (A - C) were repeated twice (n = 3) with similar results.

### Increased *PFA-DSP1* expression coincides with decreases InsP_7_ levels *in planta*

To gain insight into DSP functions *in planta*, we searched for *Arabidopsis* T-DNA insertion lines of *PFA-DSP1* and were able to identify three lines, *pfa-dsp1-3* and *pfa-dsp1-*6 in the Col-0 background and *pfa-dsp1-4* in the Ler-0 background, for which homozygous progeny could be obtained. None of these lines displayed an obvious growth phenotype under our standard growth conditions. SAX-HPLC profiles of extracts of 20-days old [^3^H]-*myo*-inositol labeled *pfa-dsp1-3* and *pfa-dsp1-4* seedlings did not reveal a significant difference compared to the respective wild-types (Figures S10A,B). However, SAX-HPLC analyses of the *pfa-dsp1-6* line revealed a significant average reduction (around 36%) of the InsP_7_/InsP_6_ ratio compared to Col-0 (Figure 6B). The levels of other InsP species remained largely unaffected (Figure 6A). The available sequencing data for this line, as well as our analysis, indicated that the insertion of the T-DNA is 18 bp upstream of the start codon, suggesting that the full-length transcript and DSP1 protein might be expressed in this line. We therefore conducted qPCR analyses of *pfa-dsp1-6* seedlings that were grown under identical conditions as the seedlings for SAX-HPLC analyses and detected an approximately 6-fold increased expression of *PFA-DSP1* in *pfa-dsp1-6* in comparison to Col-0 seedlings (Figure 6C).

**Figure 6:**
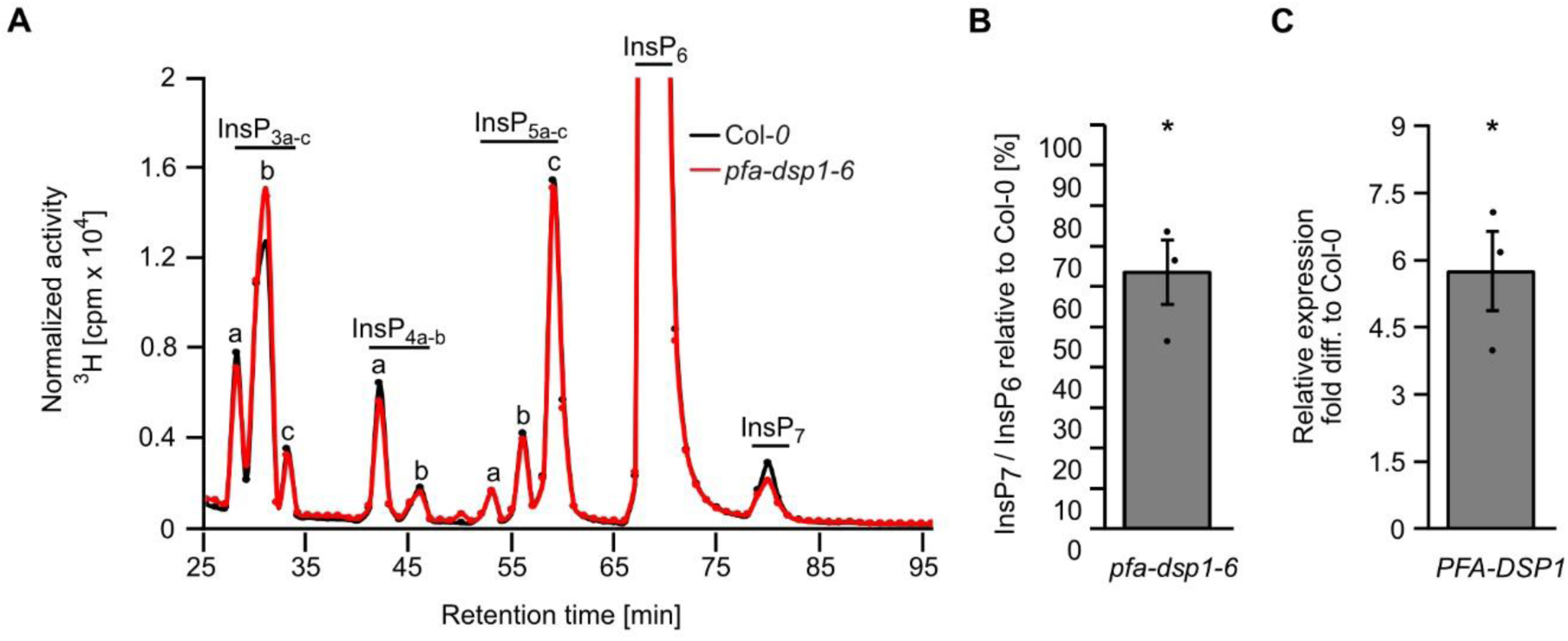
Increased expression of *PFA-DSP1* in *Arabidopsis* decreases InsP_7_ levels. (A) A representative SAX-HPLC profile of 20-days-old wild-type (Col-0) and *pfa-dsp1-6 Arabidopsis* seedlings radiolabeled with [^3^H]-*myo*-inositol. All visible peaks are highlighted and assigned to the corresponding InsP species. Based on published chromatographic mobilities ^*52, 53*^, InsP_4a_ likely represents Ins(1,4,5,6)P_4_ or Ins(3,4,5,6)P_4_, InsP_5a_ likely represents InsP_5_ [2-OH], InsP_5b_ likely represents InsP_5_ [4-OH] or its enantiomeric form InsP_5_ [6-OH], and InsP_5c_ likely represents InsP_5_ [1-OH] or its enantiomeric form InsP_5_ [3-OH]. The isomeric natures of InsP_3a-c_, InsP_4b_, InsP_7_, and InsP_8_ are unknown. (B) InsP_7_/InsP_6_ ratio of *dsp1-6* relative to Col-0 determined by analysis of SAX-HPLC profiles using OriginPro 8. (C) Relative expression of *PFA-DSP1* in plants grown under identical conditions as for SAX-HPLC analyses, presented as fold difference compared to Col-0. (B, C) Data represent means ± SEM (n = 3). Asterisks indicate values that are significantly different from the control (according to Student’s *t* test, P<0.05 (*); P<0.01 (**); P<0.001 (***)).

Since the analyses of the *pfa-dsp1-6* line indicated that the T-DNA insertion causes an overexpression of *PFA-DSP1*, resulting in decreased InsP_7_ levels, we investigated whether PP-InsP phosphohydrolase activity is also observed in a heterologous plant expression system. To this end, we transiently expressed a translational fusion of PFA-DSP1 with a C-terminal EYFP under control of the strong viral CaMV *35S* promoter in *Nicotiana benthamiana* using agrobacterium-mediated transfection. The respective catalytically inactive PFA-DSP1^C150S^-EYFP fusion protein was also expressed and InsPs were then extracted from *Nicotiana benthamiana* leaves and purified by TiO_2_ pulldown 5 days after infiltration. PAGE analyses showed that transient expression of DSP1 or expression of its catalytic inactive version did not alter InsP_6_ levels (Figure 7A). In contrast, InsP_7_ levels were reduced by the transient expression of PFA-DSP1 but not by expression of its catalytic inactive version (Figure 7A). These findings were strengthened by subsequent CE-ESI-MS analyses that revealed no changes in the ratios of 1/3-InsP_7_/InsP_6_ or 4/6-InsP_7_/InsP_6_ as compared to control leaves infiltrated with agrobacteria carrying the silencing inhibitor *P19* alone (Figure 7B). In contrast, the 5-InsP_7_/InsP_6_ ratio was significantly reduced in plants expressing *PFA-DSP1* compared to plants expressing the inactive version of *PFA-DSP1* or *P19* alone. The InsP_8_/InsP_6_ ratio, in turn, was strongly reduced by the expression of *PFA-DSP1* in agreement with a partial hydrolytic activity of DSP proteins against InsP_8_ isomers (Figures 3 and S7) and in agreement with the finding that 5-InsP_7_, a substrate hydrolyzed by PFA-DSP1, represents the major precursor for InsP_8_ synthesis ^*11*^. In summary, these results demonstrate that ectopic expression of *Arabidopsis PFA-DSP1* results in a specific decrease of 5-InsP_7_ and InsP_8_ *in planta*.

**Figure 7:**
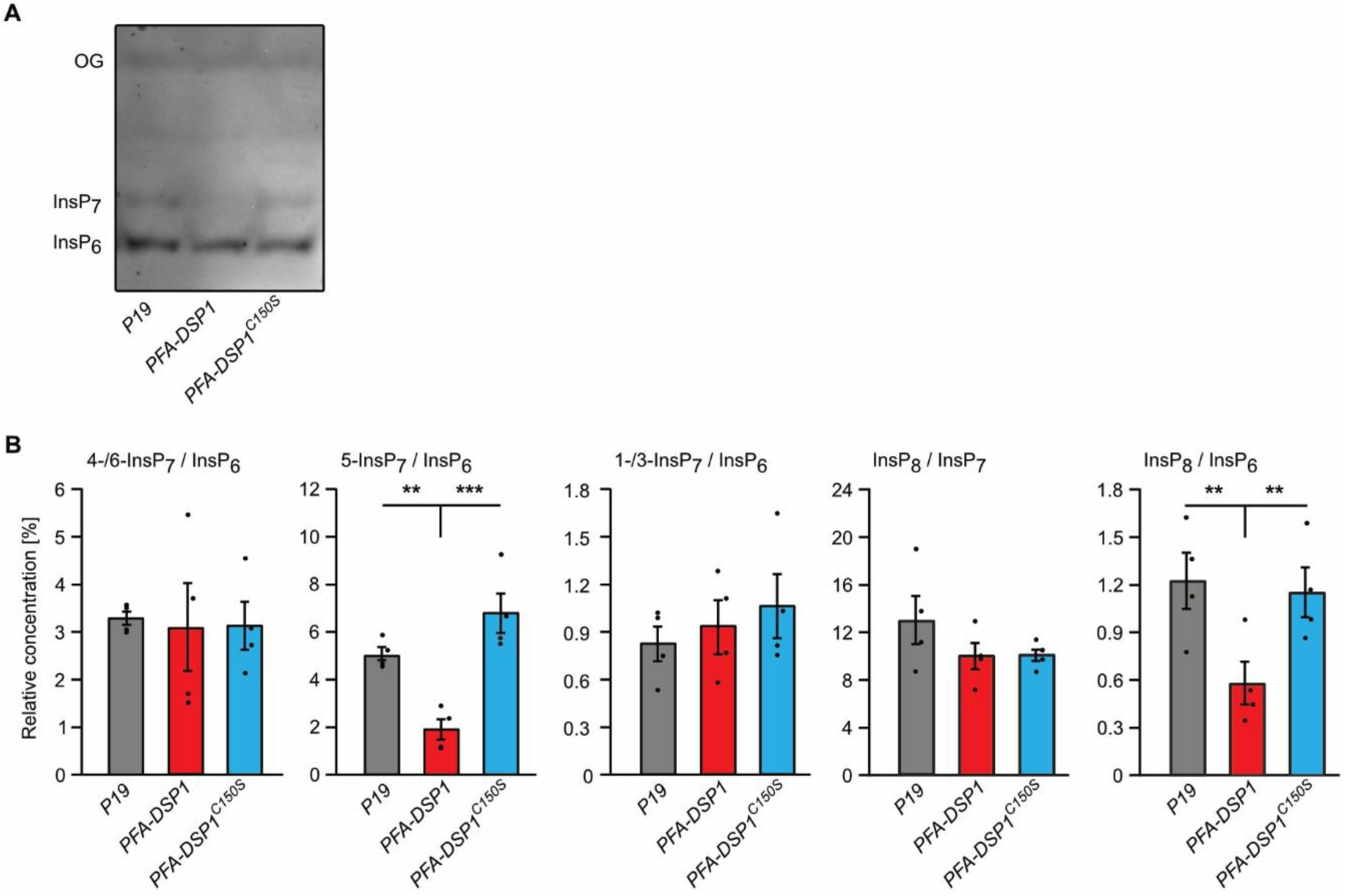
Transient expression of *PFA-DSP1* in *Nicotiana benthamiana* leaves specifically decreases 5-InsP_7_ and InsP_8_. InsPs from infiltrated *Nicotiana benthamiana* leaves, transiently expressing the silencing inhibitor *P19* alone or together with *DSP1-EYFP* or *DSP1*^*C150S*^*-EFYP*, were purified with TiO_2_ pulldown and analyzed using CE-ESI-MS. Detected peaks were assigned to specific InsP isomers and quantified by comparing them with ^13^C InsP standards that were spiked into the purified samples. (A) A representative PAGE gel of a sample set used for CE-ESI-MS analysis. (B) Relative amounts of PP-InsPs compared to InsP_6_ or of InsP_8_ compared to all InsP_7_ isomers as indicated. Data represent means ± SEM (n = 4). Asterisks indicate values that are significantly different from wild-type (according to Student’s *t* test, P<0.05 (*); P<0.01 (**); P<0.001 (***)). Note that 4- or 6-InsP_7_, as well as 1- or 3-InsP_7_ represent enantiomeric forms that cannot be distinguished by CE-ESI-MS analyses.

## Conclusions

Recent studies elucidating the identity and substrate specificity of InsP_6_/PP-InsP kinases have allowed to establish important functions of PP-InsPs in nutrient sensing, hormone signaling and plant immunity ^*5-13, 20, 28*^. In contrast, information on enzymatic activities removing PP-InsPs to switch off their signaling functions in plants is sparse. Intriguingly, the first robust detection of PP-InsP messengers in mammalian cells was made possible by blocking mammalian PP-InsP phosphohydrolases with fluoride ^*1*^. While substantial progress in elucidating the role of various PP-InsP phosphohydrolases in regulating these messengers in yeast and mammalian cells has been made ^*37, 41, 42, 54*^, we are unaware of any study about PP-InsP degrading enzymes in plants on the onset of this study. Here, we provide evidence that the *Arabidopsis* PFA-DSP proteins are functional homologs of yeast Siw14 with high phosphohydrolase specificity for the 5-β-phosphate of various PP-InsPs.

The striking biochemical similarities between *Arabidopsis PFA-DSPs* as deduced from *in vitro* assays and heterologous expressions analyses in yeast might well explain redundancies of these enzymes and consequently a lack of obvious phenotypes in single *pfa-dsp* loss-of-function lines in *Arabidopsis*. A search in transcriptome studies revealed that *PFA-DSP1, 2* and *4* are strongly induced by P_i_ deficiency (Figure S11). Such P_i_-dependent regulation is in line with the disappearance of PP-InsPs in tissues of P_i_-starved plants (Dong et al., 2019; Riemer et al., 2021), but future studies are required to establish the involvement on PFA-DSPs in the removal of messengers controlling P_i_ signaling. The high specificity of PFA-DSPs observed in this study establishes these enzymes as ideal tools to investigate the physiological roles of 5-β-phosphate containing PP-InsPs in plant development, plant immunity, nutrient perception and abiotic stress tolerance. This is particularly important because of potentially confounding effects caused by the recently discovered plant 4/6-InsP_7_ ^*11*^, but also because higher order mutants involved in the synthesis of 5-β-phosphate containing PP-InsPs such as *itpk1 itpk2* and *vih1 vih2* display severe developmental defects or die at young seedling stage ^*10, 11*^. With the availability of a variety of promoters with tight spatial and temporal regulation, ectopic expression of *PFA-DSPs* in a tissue and developmentally controlled manner will provide helpful insights to unravel the roles of 5-InsP_7_ and 1/3, 5-InsP_8_ in plant development and plant physiology.

## Methods

### Plant materials and growth conditions

Seeds of *Arabidopsis thaliana* T-DNA insertion lines *pfa-dsp1-3* (WiscDsLox_473_B10, Col-0), *pfa-dsp1-4* (CSHL_GT1415, Ler-0) and *pfa-dsp1-6* (SAIL_116_C12, Col-0) were obtained from The European Arabidopsis Stock Centre (http://arabidopsis.info/). In order to identify homozygous lines, F2 and F3 plants were genotyped by PCR using the primers indicated in Table S1.

For sterile cultures, *Arabidopsis* seeds were surface sterilized in 1.2 % (v/v) NaHClO_4_ and 0.05 % (v/v) Triton X-100 for 3 min, in 70 % (v/v) ethanol and 0.05 % (v/v) Triton X-100 for 3 min and in 100 % (v/v) ethanol before transferring onto sterile filter paper. Sterilized seeds were sown onto half-strength Murashige and Skoog (MS) medium ^*55*^ containing 1 % sucrose, pH 5.7 and solidified with 0.7 % (w/v) Phytagel (Sigma-Aldrich). After 2 days of stratification at 4°C, the plates were transferred to a growth incubator and the seedlings were grown under short-day conditions with the following regime: 8/16 h light/dark; light intensity 120 μmol m^-2^ s^-1^; temperature 22 °C/20 °C.

### Constructs

The following full-length ORFs were amplified by PCR from an *Arabidopsis* whole seedling cDNA preparation: *PFA-DSP1* (At1g05000), *PFA-DSP2* (At2g32960), *PFA-DSP3* (At3g02800) *PFA-DSP4* (At4g03960), and *PFA-DSP5* (At5g16480). Likewise, the *SIW14* ORF sequence was amplified from yeast genomic DNA. Primers used for amplification are listed in Table S1. The reverse primers contained a *V5* sequence (underlined) allowing a translational fusion of the resulting gene products with a C-terminal V5 epitope tag. Amplification products were cloned into pDONR221 (Invitrogen) via BP clonase II (Invitrogen) reaction following the manufacturer’s instructions. The ORFs were then swapped into the episomal yeast expression vector pDRf1-GW ^*56*^ by the LR clonase II (Invitrogen) reaction following the manufacturer’s instructions. For expression of *SIW14* under control of the endogenous promoter from a *CEN*-based plasmid, the *SIW14* gDNA was amplified from purified yeast gDNA using primers listed in Table S1. The *SIW14* gDNA was inserted into YCplac33 (ATCC #87586) using the restriction enzymes *Pst*I and *Eco*RI.

For protein expression, *PFA-DSP1 – 5* were amplified as described before but with a reverse primer containing a stop codon. Amplified products were cloned into pDONR221 (Invitrogen), then swapped by LR clonase II (Invitrogen) into the bacterial expression vector pDEST566 (Addgene plasmid # 11517), which contains a sequence encoding an N-terminal His_6_-maltose binding protein (MBP) epitope tag. Free His-tagged MBP protein was expressed from a modified pET28 vector carrying an N-terminal sequence encoding a His_8_-maltose binding protein (MBP) epitope tag ^*5*^.

For transient expression in *Nicotiana benthamiana*, the ORF of *PFA-DSP1* (wild-type sequence and with a mutated sequence encoding the C150S substitution) was swapped by LR clonase II (Invitrogen) from pDONR221 into the plant expression vector pGWB641 ^*57*^, which harbors a viral CaMV *35S* promoter to allow gene expression and a sequence encoding a C-terminal EYFP tag. Site-directed mutagenesis was performed on the respective plasmids with primers listed in Table S1.

### *Nicotiana benthamiana* infiltration

A single colony of transformed *Agrobacteria* was inoculated in 2 mL LB media containing the appropriate antibiotics and cultivated overnight at 26°C in a spinning wheel. On the next morning, 1 mL of overnight culture was added to 5 mL fresh LB with antibiotics and grown for another 4 h at 26°C. Afterwards, the cultures were harvested by centrifugation at 4°C with 3000 *g* for 20 min. The pellet was then resuspended in 3 mL infiltration solution containing 10 mM MgCl_2_, 10 mM MES-KOH (pH 5.6) and 150 μM acetosyringone. OD_600_ was determined using a 1:10 dilution and adjusted to 0.8 in infiltration solution. Then the working solution was prepared by pooling equal amounts of cultures that were co-infiltrated (e.g. *P19* + *PFA-DSP1*). The abaxial surface of the leaf (lower side) was then infiltrated using a 1 mL syringe without a needle. Afterwards, the plants were placed in a dark incubator at 26°C for approx. 1 day before keeping them for another 4 days on the workbench. The leaves were then harvested and frozen in liquid nitrogen before continuing with the extraction of inositol phosphates.

### Yeast Strains

Different strains of the budding yeast *Saccharomyces cerevisiae* were used. The BY4741 wild-type (MATa *his3Δ leu2Δ met15Δ ura3Δ*), *siw14Δ* (*YNL032w::kanMX4*), *vip1Δ* (*YLR410w::kanMX4*), *kcs1Δ* (*YDR017c::kanMX4*), and *ipk2Δ* (*YDR173c::kanMX4*) were obtained from Euroscarf. *vip1Δ siw14Δ, kcs1Δ siw14Δ, ipk2Δ siw14Δ* were generated by using *loxP*/Cre gene disruption and the *ble* resistance marker, which confers phleomycine/Zeocin (Invitrogen) resistance ^*58*^ using primers listed in Table S1. In addition, the following mutants in the DDY1810 background (MATa; *leu2Δ ura3-52 trp1Δ; prb1-1122 pep4-3 pre1-451*) ^*59*^ were used: *kcs1Δ* and *kcs1Δ ddp1Δ. kcs1Δ siw14Δ, kcs1Δ ddp1Δ siw14Δ* and *siw14Δ* were generated in this background as described before. For all assays, the yeast cells were transformed by the Li-acetate method ^*60*^ and cultured either in 2 x YPD + CSM medium or selective synthetic deficiency (SD) medium.

### Yeast growth complementation assay

Yeast transformants were inoculated in selective synthetic deficiency (SD) medium and grown overnight at 28°C while shaking (200 rpm). Then the OD_600_ was measured, adjusted to 1.0 and an 8-fold dilution series was prepared in a 96-well plate. Subsequently, 10 μL of each dilution were spotted on selective solid media as described earlier ^*61*^ and incubated at 26°C for 2 - 4 days. To prepare selective solid media supplemented with wortmannin, autoclaved media was cooled down to 60°C, wortmannin was added from a 10 mM stock in DMSO (Sigma Aldrich) to a final concentration of 1 - 3 μM. Since the activity of wortmannin changed by age and by the number of freezing/thawing cycles, aliquots were kept at −20°C and were not thawed more than five times. In addition, several concentrations were employed for the spotting assays to be able to identify the activity at which growth differences between *siw14Δ, kcs1Δ* and their isogenic wildtype transformants became most obvious. Pictures were taken with a Bio-Rad ChemiDoc MP imager using white backlight.

### Protein preparation

His_6_-MBP-PFA-DSP protein fusions or free His_8_-MBP were expressed in *Escherichia coli* BL21 CodonPlus (DE3)-RIL cells (Stratagene). Overnight bacterial cultures were inoculated 1:1000 into fresh 2YT medium (1.6 % tryptone, 1 % yeast extract, 0.5 % NaCl) with 100 mg/L ampicillin (pDEST566) or 50 mg/L kanamycin (pET28) and 25 mg/L chloramphenicol. Cells were grown at 37°C while shaking (200 rpm) for 4 h (approx. 0.6 OD_600_), and protein expression was induced at 16°C overnight with 0.1 mM isopropyl-D-1-thiogalactopyranoside. Cells were lysed as described ^*62*^ using the following lysis buffer: 50 mM Tris-Cl, pH 7.5, 100 mM NaCl, 25 mM imidazole, 10 % (v/v) glycerol, 0.1 % (v/v) Tween 20, 5 mM β-mercaptoethanol and the EDTA-free complete ULTRA protease inhibitor cocktail (Roche). Proteins were batch-purified using Ni-NTA agarose resin (Macherey-Nagel) and eluted using the abovementioned lysis buffer with increased imidazole concentration (250 mM). Three elutions were combined and dialyzed using Slide-A-Lyzer™ Dialysis Cassettes (Thermo Scientific) following the manufacturer’s instructions and a buffer containing 50 mM Tris-Cl, pH 7.5 and 100 mM NaCl. The concentrated protein preparations were then stored at −20 °C. Purified proteins were analyzed using SDS-PAGE followed by Coomassie blue staining. Proteins were compared with PageRuler plus prestained protein ladder (Thermo Fisher) and with designated amounts of a BSA standard to estimate target protein concentrations.

### *In vitro* PP-InsP phosphohydrolase assay

The phosphohydrolase assay was carried out in a 15 μl reaction mixture containing 0.35 μM – 2 μM recombinant PFA-DSP or Siw14 protein, 50 mM HEPES (pH 7.0), 10 mM NaCl, 5 % (v/v) glycerol, 0.1 % (v/v) β-mercaptoethanol and 0.33 mM of various InsP_7_ and InsP_8_ isomers as indicated, and was incubated for 1 h, 2 h or 24 h at 22°C. The PP-InsP isomers were synthesized as described previously ^*63, 64*^. Reactions were separated by 33% PAGE and visualized by toluidine blue or DAPI staining.

### Titanium dioxide bead extraction and PAGE/CE-ESI-MS

Purification of inositol polyphosphates using TiO_2_ beads and analysis via PAGE was performed as described previously ^*11*^. CE-ESI-MS analyses of *in vitro*, yeast and plant samples were performed as described previously ^*11, 23*^.

### Inositol polyphosphate extraction from yeast cells and seedlings and HPLC analyses

For inositol polyphosphate analyses from yeast, transformants were inoculated into selective synthetic deficiency (SD) medium and grown overnight at 28 °C while shaking (200 rpm). They were then diluted 1:200 in 2 mL fresh medium supplemented with 6 μCi mL^−1^ [^3^H]-*myo*-inositol (30–80 Ci mmol^−1^; Biotrend; ART-0261-5) and grown overnight at 28°C in a spinning wheel. After harvest of the cell pellet, inositol polyphosphates were extracted and analysed as described before ^*5, 65, 66*^.

Extraction of [^3^H]-*myo*-inositol polyphosphates from *Arabidopsis* seedlings and subsequent SAX-HPLC analyses were performed as described previously ^*66*^.

### RNA isolation and quantitative real-time PCR

15-days old seedlings were transferred from solid half-strength MS plates to liquid half-strength MS media (supplemented with 1 % sucrose) for 5 days before harvest and immediately frozen in liquid N_2_. Total RNA was extracted with NuceloSpin RNA Plant and Fungi kit (Macherey-Nagel). cDNA was synthesized using RevertAid RT reverse transcription kit (Thermo Fisher). Quantitative PCR reactions were conducted with the CFX384 real-time system (Bio-Rad) and the SsoAdvanced Universal SYBR Green Supermix (Bio-Rad) using the primers listed in Table S1. *TIP41like* and *PP2AA3* were used as reference genes to normalize relative expression levels of all tested genes. Relative expression was calculated using the CFX Maestro software (Bio-Rad).

### Yeast protein extraction and immunodetection

Multiple transformants were inoculated into 4 mL of YPD (with 3 % glucose) or selective SD-media and grown for up to 24 h at 28°C. On the following day, the yeast was re-inoculated into 4 mL fresh media and grown for another day. Afterwards the cells were harvested and resuspended in 500 μL extraction buffer (300 mM sorbitol, 150 mM NaCl, 50 mM Na_2_HPO_4_, 1 mM EDTA, pH 7.5), supplemented with 100 mM β-mercaptoethanol and a 1:50 dilution of protease inhibitor cocktail for fungal extracts (Sigma-Aldrich). Cells were lysed with bead beating using 150 – 200 μL of glass beads (ø 0.5 mm). Lysate was spun down and the supernatant boiled for 10 min after addition of sample buffer. The protein extracts were then resolved by SDS-PAGE. Target proteins were detected by immunoblot. As primary antibody a mouse anti-V5 (Invitrogen, R960-25, 1:2000 dilution) antibody was used, followed by an Alexa fluor plus 800 goat anti-mouse antibody (Invitrogen, 1:20000 dilution). As loading control, Gal4 was detected using a rabbit polyclonal anti-Gal4 antibody (Santa Cruz, 1:1000 dilution), followed by a goat anti-rabbit StarBright Blue 700 antibody (Bio-Rad, 1:2500 dilution). The signal was detected using the multi-plex function of the ChemiDoc MP imager (Bio-Rad)

### Accession numbers

DNA and Protein Sequences can be obtained from the Saccharomyces Genome database (https://www.yeastgenome.org/), TAIR (https://www.arabidopsis.org) and UniProt (https://www.uniprot.org/) under the following accession numbers: *SIW14* (YNL032W, NC_001146.8), *Arabidopsis PFA-DSP1* (At1g05000, NM_100379.3), *Arabidopsis PFA-DSP2* (At2g32960, NM_128856.5), *Arabidopsis PFA-DSP3* (At3g02800, NM_111148.3), *Arabidopsis PFA-DSP4* (At4g03960, NM_116634.4), *Arabidopsis PFA-DSP5* (At5g16480, NM_121653.4), *Arabidopsis PP2AA3* (At1g13320, NM_101203) and *Arabidopsis TIP41-like* (At3g54000, NM_115260).

## Supporting Information

The following supporting information is available.

**Figure S1:** Purification of PFA-DSP proteins.

**Figure S2:** *In vitro, Arabidopsis* PFA-DSP1 displays robust PP-InsP phosphohydrolase activity against 5-InsP_7_ and partial phosphohydrolase activity against 4-InsP_7_ and 6-InsP_7_, respectively. **Figure S3:** In the absence of divalent cations, all InsP_7_ isomers with the exception of 2-InsP_7_ become substrates for selected *Arabidopsis* PFA-DSPs *in vitro*.

**Figure S4:** In the presence of Mg^2+^, PFA-DSP1 and PFA-DSP3 display robust *in vitro* InsP_7_ phosphohydrolase activity with high specificity for the 5-β-phosphate.

**Figure S5:** Under prolonged incubation time, *Arabidopsis* PFA-DSP1 efficiently hydrolyzes 5-InsP_7_, 4-InsP_7_ and 6-InsP_7_ but only displays partial activities against 1-InsP_7_ and 3-InsP_7_, and a very weak activity against 2-InsP_7_.

**Figure S6:** *Arabidopsis* PFA-DSP1 maintains 5-InsP_7_ phosphohydrolase activity during prolonged incubation times *in vitro*.

**Figure S7:** *In vitro, Arabidopsis* PFA-DSPs display robust 1/3,5-InsP_8_ phosphohydrolase activity.

**Figure S8:** Heterologous expression of *Arabidopsis* PFA-DSPs complements *siw14Δ*-associated defects in InsP_7_/InsP_6_ ratios in yeast.

**Figure S9:** Complementation of *siw14Δ*-associated growth defects depends on catalytic activity.

**Figure S10:** Single mutant *Arabidopsis pfa-dsp1* loss-of-function lines do not display InsP/PP-InsP defects.

**Figure S11:** *Arabidopsis* PFA-DSP1, 2 and 4 are strongly induced by P_i_ deficiency.

**Table S1:** Oligonucleotide sequences.

## Author Contributions

D.L., G.S., and P.G. conceived the study. P.G., D.L., G.S., H.J.J., and R.F.H.G. designed experiments. P.G., R.S., D.L., G.L. D.Q., J.W., M.H., N.J., K.R., J.S., N.F.R., R.F.H.G. performed experiments. P.G. generated yeast mutants and performed all yeast experiments, generated constructs, isolated T-DNA insertion lines, performed HPLC analyses of plants, performed qPCR analyses, performed plant infiltration and TiO_2_ pulldowns and analysed most of the experiments. R.S. purified recombinant proteins and carried out and analysed *in vitro* kinase assays. G.L. and D.Q. performed CE-ESI-MS/MS analysis and isomer identification. J.W. and J.S. generated constructs and established the expression and purification of recombinant proteins. N. J., M.H., and K.R. synthesized InsP_7_ and InsP_8_ isomers. N.F.R. isolated T-DNA insertion lines, performed HPLC analyses of plants, generated constructs and performed qPCR analyses. R.F.H.G. generated plant samples for CE-ESI-MS analysis and did transcriptome analysis. M.N.T. synthesized ^13^C-InsP standards. V.G. analysed and quantified HPLC analyses. P.G., G.S., D.L., H.J.J. and D.F. supervised the experimental work. P.G., G.S., R.S., D.L., and R.Y. wrote the manuscript with input from all authors.

## Notes

The authors declare no competing financial interest.

## Acknowledgments

We thank Li Schlüter and Brigitte Ueberbach (Department of Plant Nutrition, Institute of Crop Science and Resource Conservation, University of Bonn) for excellent technical assistance and Marília Kamleitner (Department of Plant Nutrition, Institute of Crop Science and Resource Conservation, University of Bonn) for critically reading this manuscript.

## Funding Sources

This work was funded by grants from the Deutsche Forschungsgemeinschaft (SCHA 1274/4-1, SCHA 1274/5-1, Research Training Group GRK 2064 and Germany’s Excellence Strategy, EXC-2070-390732324, PhenoRob to G.S.; JE 572/ 4-1 and Germany’s Excellence Strategy, CIBSS–EXC-2189–Project ID 390939984 to H.J.J; and HE 8362/1-1, DFG Eigene Stelle, to R.F.H.G.). D.L acknowledges the Department of Biotechnology (DBT) for HGK-IYBA award (BT/13/IYBA/2020/04) and a DBT-IISc Partnership Program. H.J.J. acknowledges funding from the Volkswagen Foundation (Momentum Grant 2021)

^&^Present address: Department of Biomedicine, University of Basel, 4058 Basel, Switzerland

