## supporting information for "*Arabidopsis* PFA-DSP-type phosphohydrolases target specific inositol pyrophosphate messengers"

#### Short title

***A. thaliana* inositol pyrophosphate phosphohydrolases**

#### Article title

### *Arabidopsis* PFA-DSP-type phosphohydrolases target specific inositol pyrophosphate messengers

Philipp Gaugler[1§], Robin Schneider[1§], Guizhen Liu[2], Danye Qiu[2], Jonathan Weber[1], Jochen Schmid[3&], Nikolaus Jork[2,4], Markus Häner[2], Kevin Ritter[2], Nicolás Fernández-Rebollo[3], Ricardo F.H. Giehl[5], Minh Nguyen Trung[6], Ranjana Yadav[7], Dorothea Fiedler[6], Verena Gaugler[1], Henning J. Jessen[2], Gabriel Schaaf[1]*, Debabrata Laha[7]*

[1]Department of Plant Nutrition, Institute of Crop Science and Resource Conservation, Rheinische Friedrich-Wilhelms-Universität Bonn, 53115 Bonn, Germany

[2]Department of Chemistry and Pharmacy and CIBSS-Centre for Integrative Biological Signalling Studies, Albert-Ludwigs University Freiburg, 79104 Freiburg, Germany

[3]Center for Plant Molecular Biology, Department of Plant Physiology, Eberhard Karls University Tübingen, 72076 Tübingen, Germany

[4]Spemann Graduate School of Biology and Medicine (SGBM), University of Freiburg, 79104 Freiburg, Germany

[5]Department of Physiology & Cell Biology, Leibniz-Institute of Plant Genetics and Crop Plant Research, 06466 Gatersleben, Germany

[6]Leibniz-Forschungsinstitut für Molekulare Pharmakologie, 13125 Berlin, Germany; Department of Chemistry, Humboldt Universität zu Berlin, 12489 Berlin, Germany.

[7]Department of Biochemistry, Indian Institute of Science (IISc), Bengaluru-560012, India

[§]P.G. and R.S. contributed equally to this manuscript

*

ORCID IDs: P.G.: 0000-0002-9187-404X, R.S. 0000-0002-9657-4548, G.L. 0000-0003-1748-6687, DQ: 0000-0003-2197-3218, J.S.: 0000-0001-7527-7088, N.J.: 0000-0002-3717-443X, M.H.: 0000-0003-3313-4868, K.R.: 0000-0001-7900-1855, N.F.R.: 0000-0003-3556-8766, R.F.H.G: 0000-0003-1006-3163, M.N.T.: 0000-0003-3100-9609, R.Y.: 0000-0003-4956-5638, D.F.: 0000-0002-0798-946X, V.G.: 0000-0002-3249-2922, H.J.J: 0000-0002-1025-9484, GS: 0000-0001-9022-4515, D.L.: 0000-0002-7823-5489

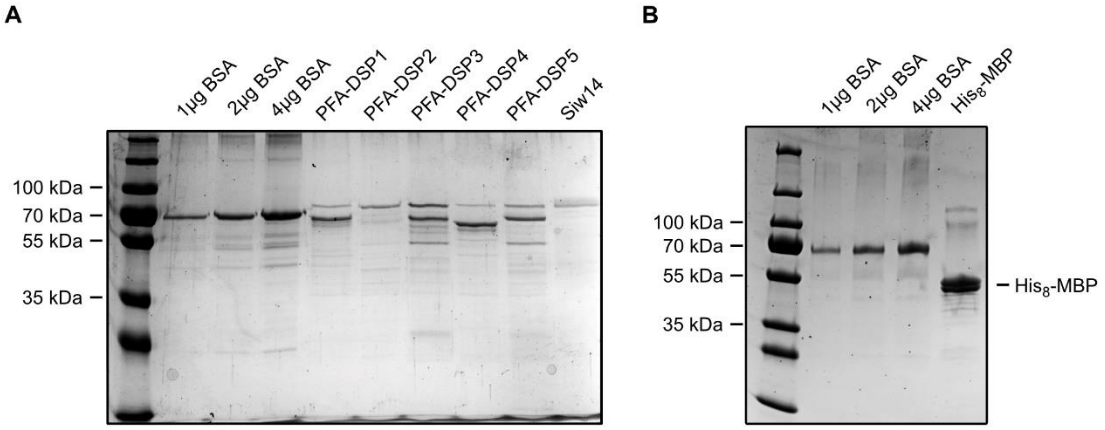


**Figure S1: Purification of PFA-DSP proteins.** (A, B) Recombinant His-MBP-DSPs or His-MBP-Siw14 were expressed in *E. coli* and purified with Ni-NTA resin as described in methods. Dialyzed proteins were denatured and separated by SDS-PAGE in parallel with BSA standards to determine protein concentrations.

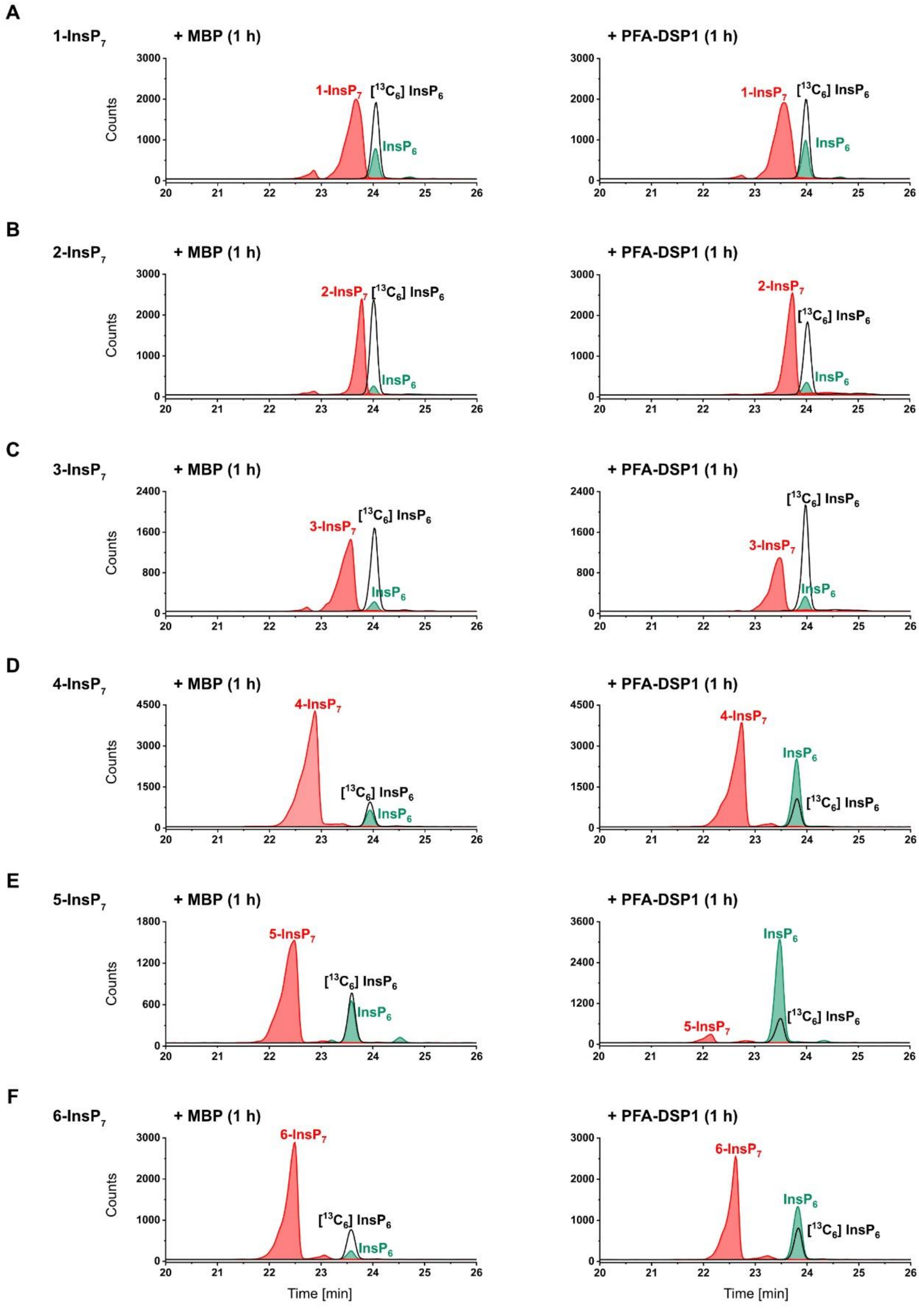


**Figure S2: *In vitro*, *Arabidopsis* PFA-DSP1 displays robust PP-InsP phosphohydrolase activity against 5-InsP$_7$ and partial phosphohydrolase activity against 4-InsP$_7$ and 6-InsP$_7$, respectively.** (A – F) 0.4 µM PFA-DSP1 was incubated with 0.33 mM InsP$_7$ and 1 mM $MgCl_2$ for 1 h. The reaction product was spiked with an isotopic standards mixture ([$^{13}C_6$]1,5-InsP$_8$, [$^{13}C_6$]5-InsP$_7$, [$^{13}C_6$]1-InsP$_7$, [$^{13}C_6$] InsP$_6$, [$^{13}C_6$]2-OH InsP$_5$) and subjected to CE-ESI-MS analyses. Representative extracted-ion electropherograms of samples shown in Figure 1.

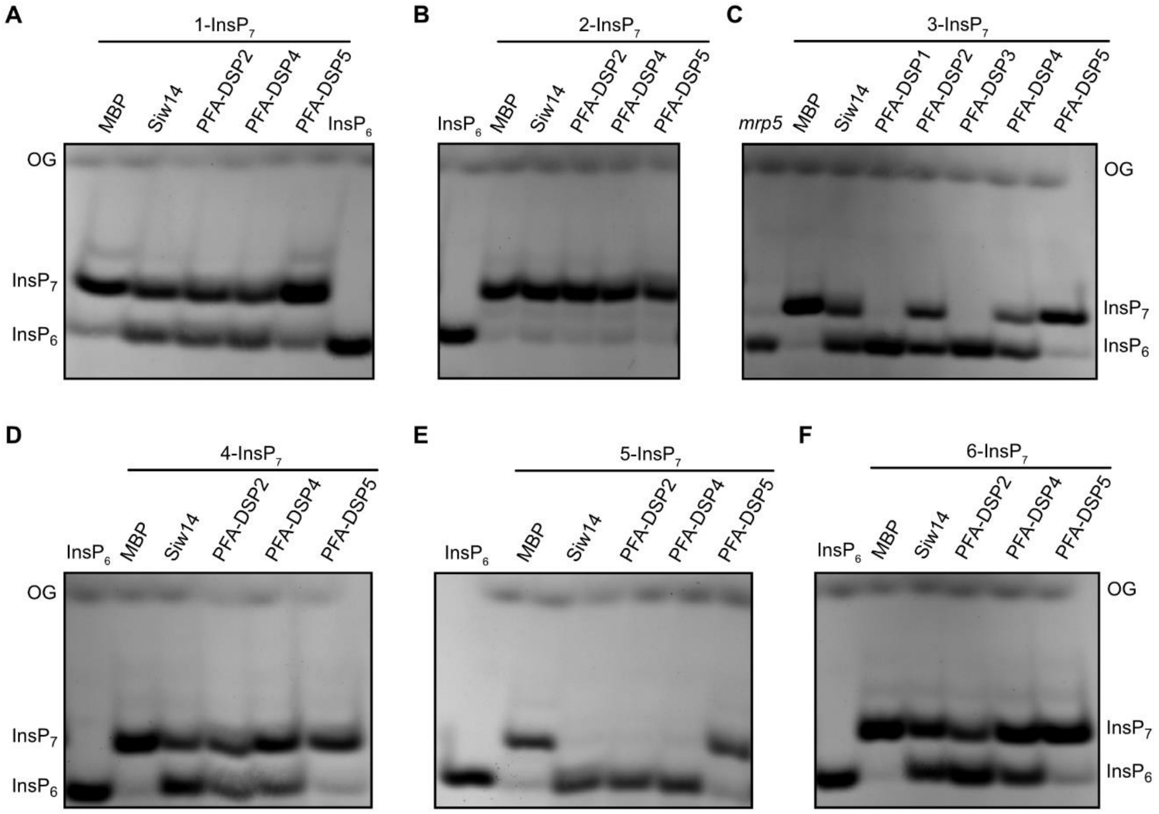


**Figure S3: In the absence of divalent cations, all $InsP_7$ isomers with the exception of 2-$InsP_7$ become substrates for selected *Arabidopsis* PFA-DSPs *in vitro*.** (A – F) Approximately 0.4 µM His-MBP-DSPs and His-MBP were incubated with 1 mM EDTA and 0.33 mM $InsP_7$ for 1 h at 22°C. His-MBP served as a negative control. The reaction products were separated by 33 % PAGE and visualized with toluidine blue.

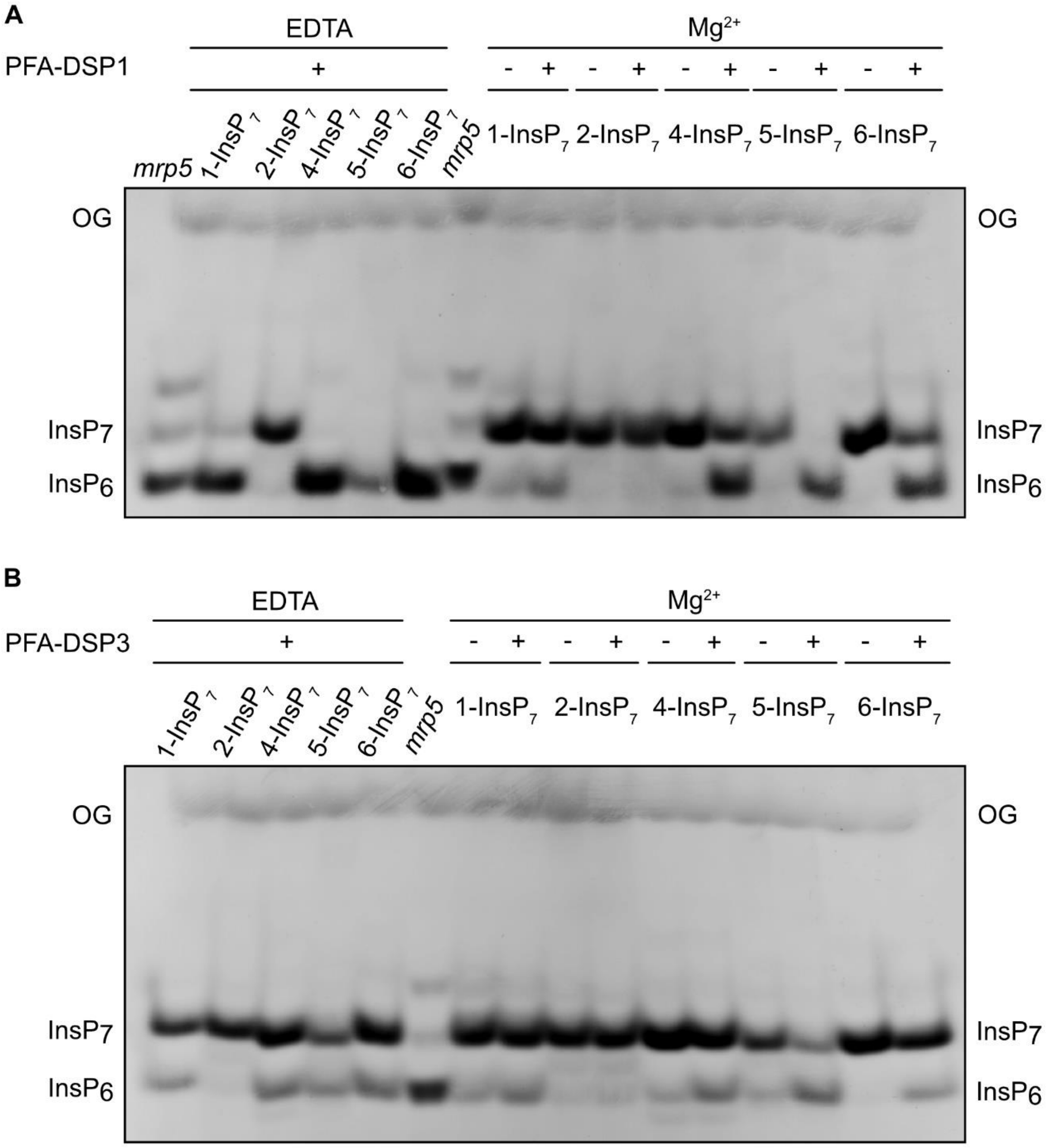


**Figure S4: In the presence of $Mg^{2+}$, PFA-DSP1 and PFA-DSP3 display robust *in vitro* $InsP_7$ phosphohydrolase activity with high specificity for the 5-β-phosphate**. (A – B) Approximately 0.4 µM His-MBP-DSP1 and His-MBP-DSP3 were incubated with 0.33 mM $InsP_7$ and 1 mM EDTA or 1 mM $MgCl_2$ for 1 h at 22°C. His-MBP served as a negative control. The reaction products were separated by 33 % PAGE and visualized with toluidine blue.

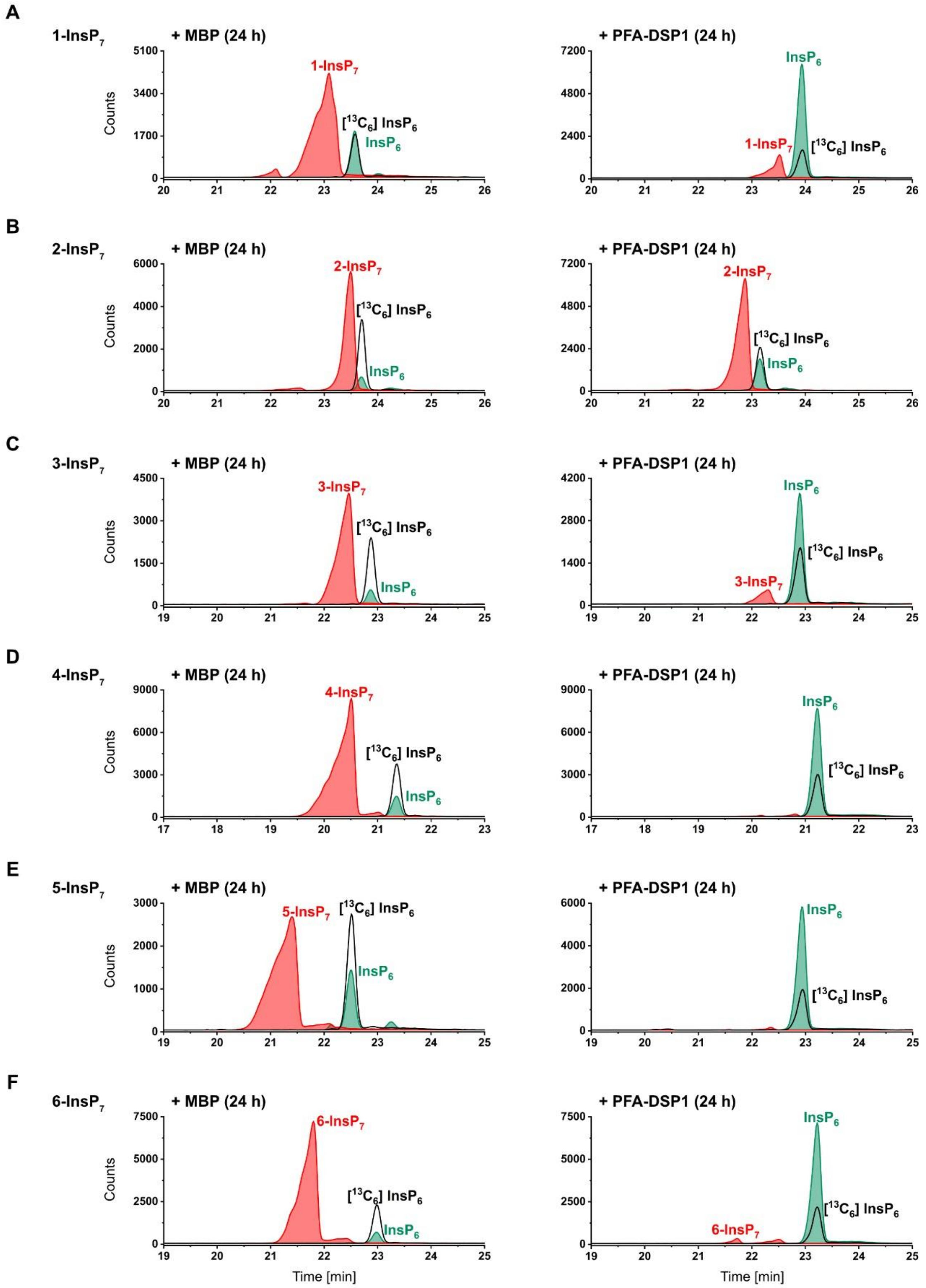


**Figure S5: Under prolonged incubation time, *Arabidopsis* PFA-DSP1 efficiently hydrolyzes 5-InsP$_7$, 4-InsP$_7$ and 6-InsP$_7$ but only displays partial activities against 1-InsP$_7$ and 3-InsP$_7$, and a very weak activity against 2-InsP$_7$.** (A – F) 0.4 µM PFA-DSP1 was incubated with 0.33 mM InsP$_7$ and 1 mM $MgCl_2$ for 24 h. The reaction product was spiked with an isotopic standards mixture ([$^{13}C_6$]1,5-InsP$_8$, [$^{13}C_6$]5-InsP$_7$, [$^{13}C_6$]1-InsP$_7$, [$^{13}C_6$] InsP$_6$, [$^{13}C_6$]2-OH InsP$_5$) and subjected to CE-ESI-MS analyses. Representative extracted-ion electropherograms of samples shown in Figure 2.

A

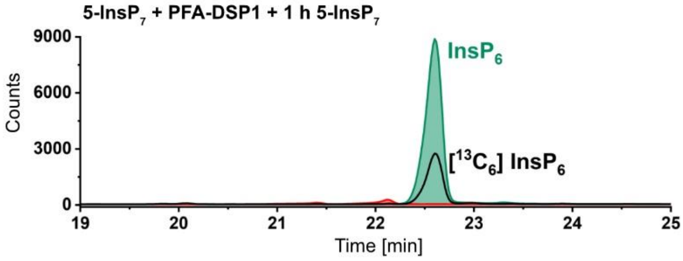


**Figure S6: *Arabidopsis* PFA-DSP1 maintains 5-InsP$_7$ phosphohydrolase activity during prolonged incubation time *in vitro*.** (A) 0.4 µM PFA-DSP1 was incubated with 0.33 mM 5-InsP$_7$ and 1 mM $MgCl_2$ for 24 h. To ensure that PFA-DSP1 is active during the whole incubation time, 0.33 mM 5-InsP$_7$ was added after 23 h and incubated for another 1 h. The reaction product was spiked with an isotopic standards mixture ([$^{13}C_6$]1,5-InsP$_8$, [$^{13}C_6$]5-InsP$_7$, [$^{13}C_6$]1-InsP$_7$, [$^{13}C_6$] InsP$_6$, [$^{13}C_6$]2-OH InsP$_5$) and subjected to CE-ESI-MS analyses.

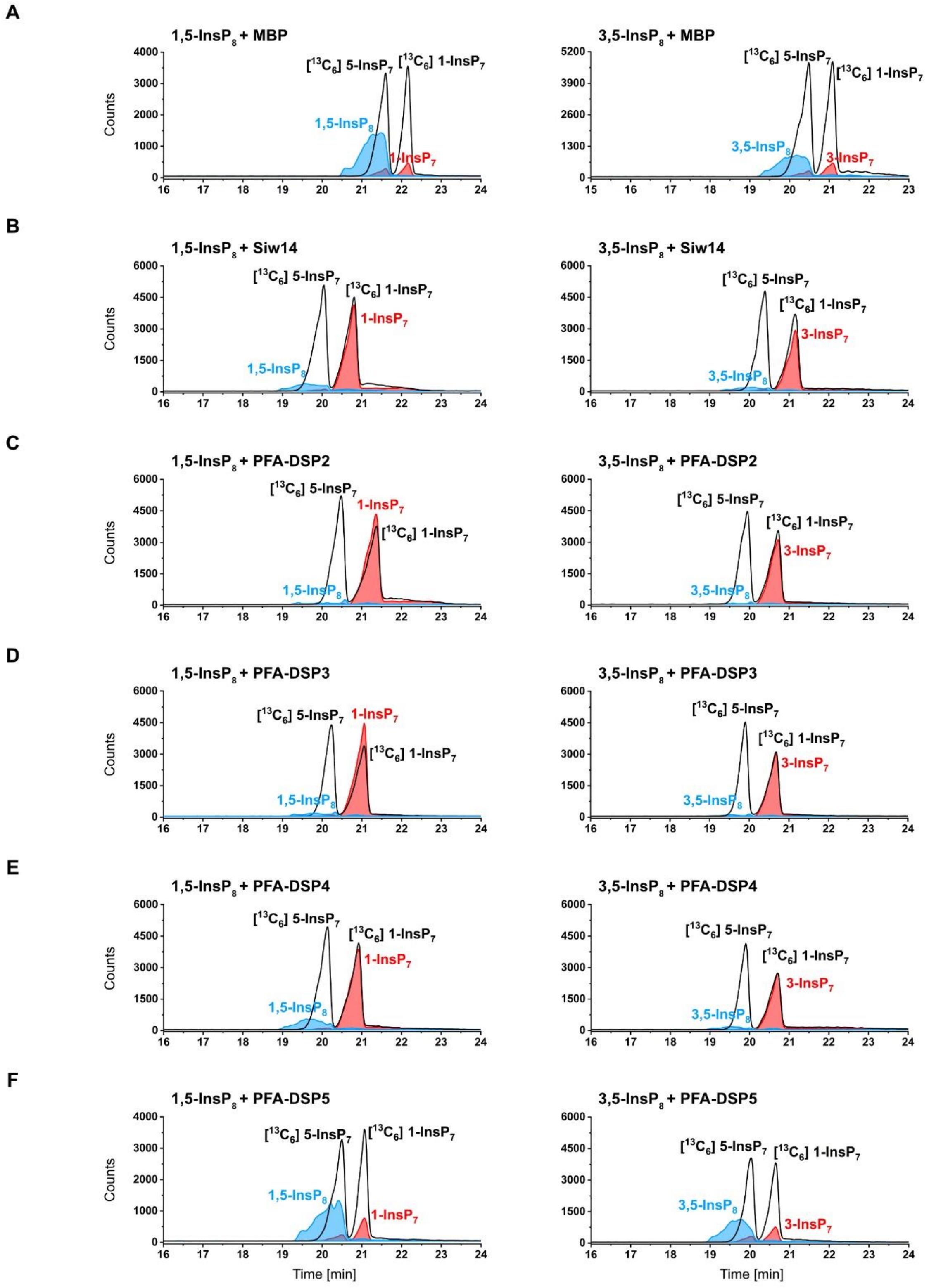


**Figure S7: *In vitro*, *Arabidopsis* PFA-DSPs display robust 1/3,5-InsP$_8$ phosphohydrolase activity.** (A – F) Approximately 0.4 µM His-MBP-DSPs and His-MBP-Siw14 were incubated with 0.33 mM 1,5-InsP$_8$ or 3,5-InsP$_8$ and 1 mM MgCl$_2$ for 1 h. The reaction products were spiked with isotopic standards mixture ([$^{13}$C$_6$]1,5-InsP$_8$, [$^{13}$C$_6$]5-InsP$_7$, [$^{13}$C$_6$]1-InsP$_7$, [$^{13}$C$_6$] InsP$_6$, [$^{13}$C$_6$]2-OH InsP$_5$) and subjected to CE-ESI-MS analyses. Representative extracted-ion electropherograms of samples shown in Figure 3C.

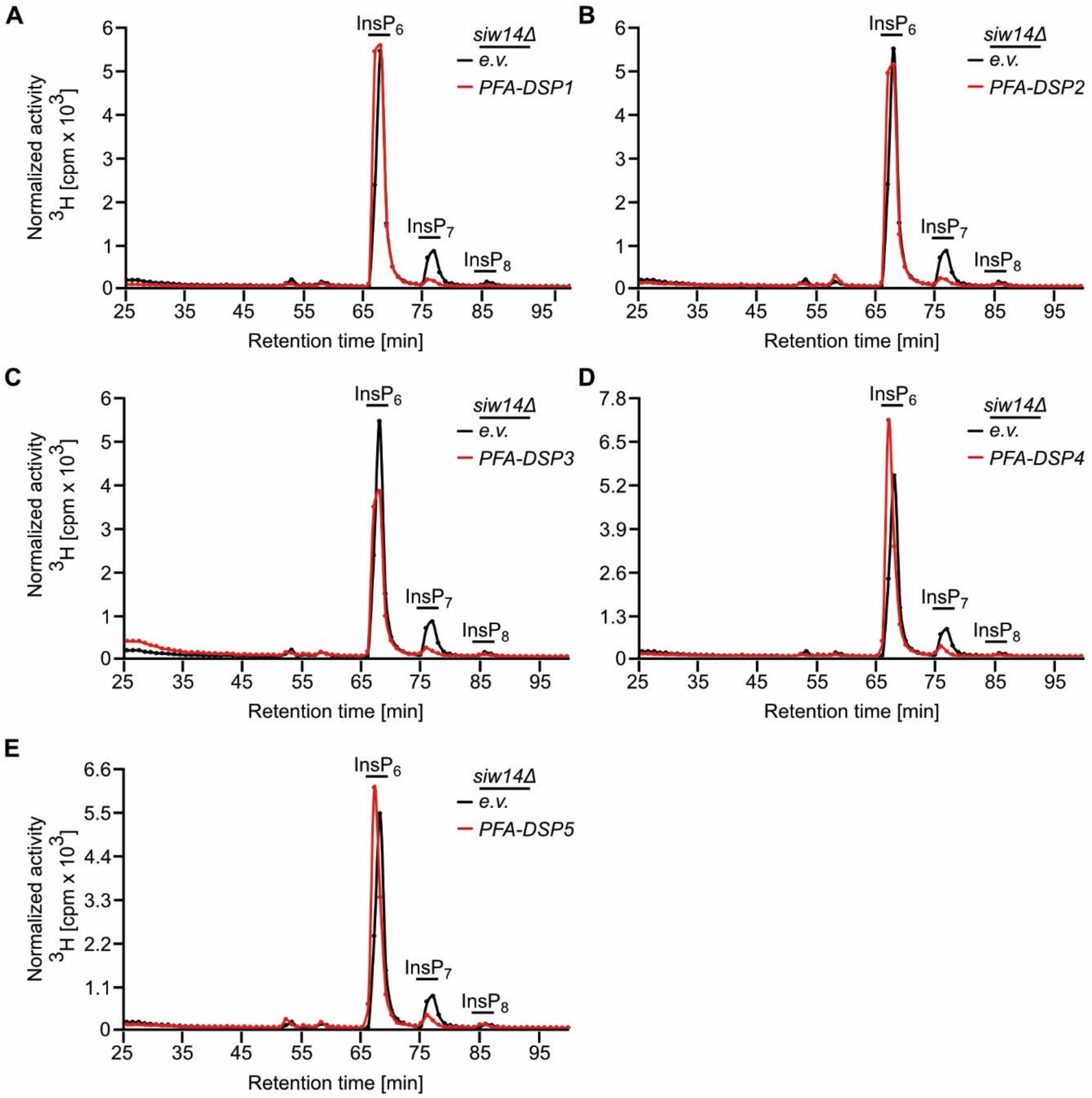


**Figure S8: Heterologous expression of *Arabidopsis* PFA-DSPs complements *siw14Δ*-associated defects in $InsP_7/InsP_6$ ratios in yeast.** (A - E) SAX-HPLC profiles of radiolabeled *siw14Δ* yeast transformed with either empty pDRf1-GW plasmid (e.v.) or pDRf1-GW carrying *PFA-DSP1 - 5*. Depicted is a representative analysis of each *PFA-DSP* transformant, with the same analysis of a representative empty vector transformant shown in the same profile in each graph. The experiment was repeated twice (n = 3) with similar results (combined data shown in Figure 4B).

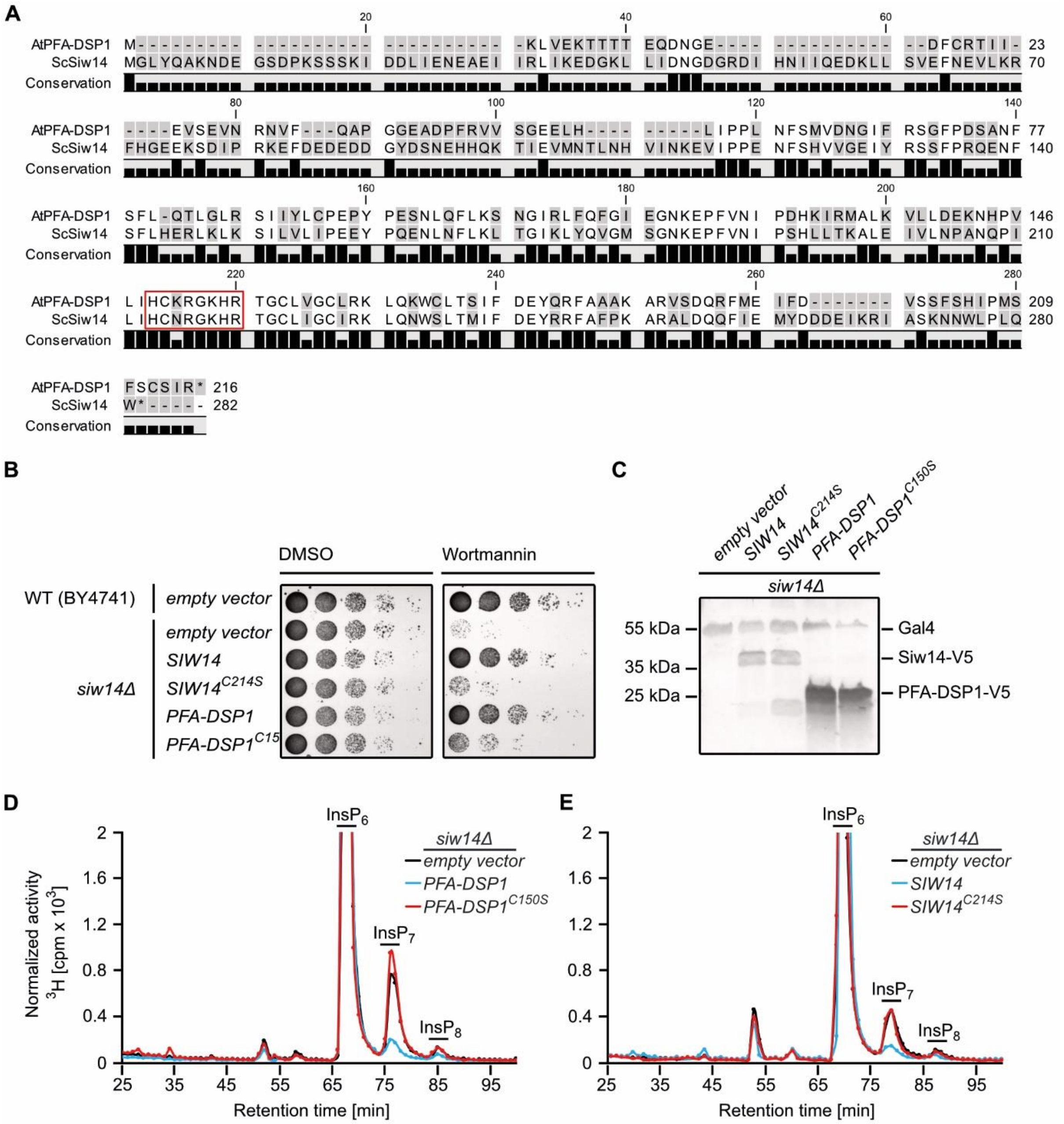


**Figure S9: Complementation of *siw14Δ*-associated growth defects depends on catalytic activity.** (A) Protein alignment of Siw14 from yeast and its homolog PFA-DSP1 from *Arabidopsis thaliana*. Identical amino acids are shown in black, different residues are highlighted with grey boxes. The conserved PTP (Protein Tyrosine Phosphatase) signature motif HC(X)5R is highlighted with the red box. The alignment was generated via the Multiple Alignments function of CLC Main Workbench 8 (QIAGEN). (B) Growth complementation assay with *siw14Δ*. Wild-type yeast (BY4741) and the *siw14Δ* yeast mutant were transformed with pDRf1-GW plasmids carrying either *SIW14* or its catalytic mutant C214S or carrying *PFA-DSP1* or its catalytic mutant C150S. Yeast strains transformed with empty pDRf1-GW vector as indicated served as controls. Transformants were then spotted in 8-fold serial dilutions (starting from OD 1.0) onto selective media containing wortmannin solved in DMSO or DMSO alone as control. Plates were incubated at 26 °C for 2 days before photographing. (C) Immunoblotting of Siw14, Siw14$^{C214S}$, PFA-DSP1 and PFA-DSP1$^{C150S}$. For detection of V5-tagged proteins an anti-V5 tag primary antibody

(Invitrogen; 1:2000 dilution) and an anti-mouse secondary antibody coupled with Alexa Fluor plus 800 (Invitrogen; goat; 1:20000 dilution) were used. As loading control, Gal4 protein levels were detected simultaneously using a polyclonal anti-Gal4 antibody (Santa Cruz; 1:1000 dilution) and an anti-rabbit StarBright Blue 700 antibody (Bio-Rad, goat; 1:2500 dilution). The signal was detected using the multiplex function of the ChemiDoc MP imager (Bio-Rad). (D) SAX-HPLC profiles of extracts of radiolabeled *siw14Δ* yeast transformed with either empty pDRf1-GW plasmid (empty vector) or pDRf1-GW carrying either *SIW14* or *SIW14*$^{C214S}$. (E) SAX-HPLC profiles of radiolabeled *siw14Δ* yeast transformed with either empty pDRf1-GW plasmid (e.v.) or pDRf1-GW carrying either *PFA-DSP1* or *PFA-DSP1*$^{C150S}$. (B – E) The experiments were repeated independently with similar results.

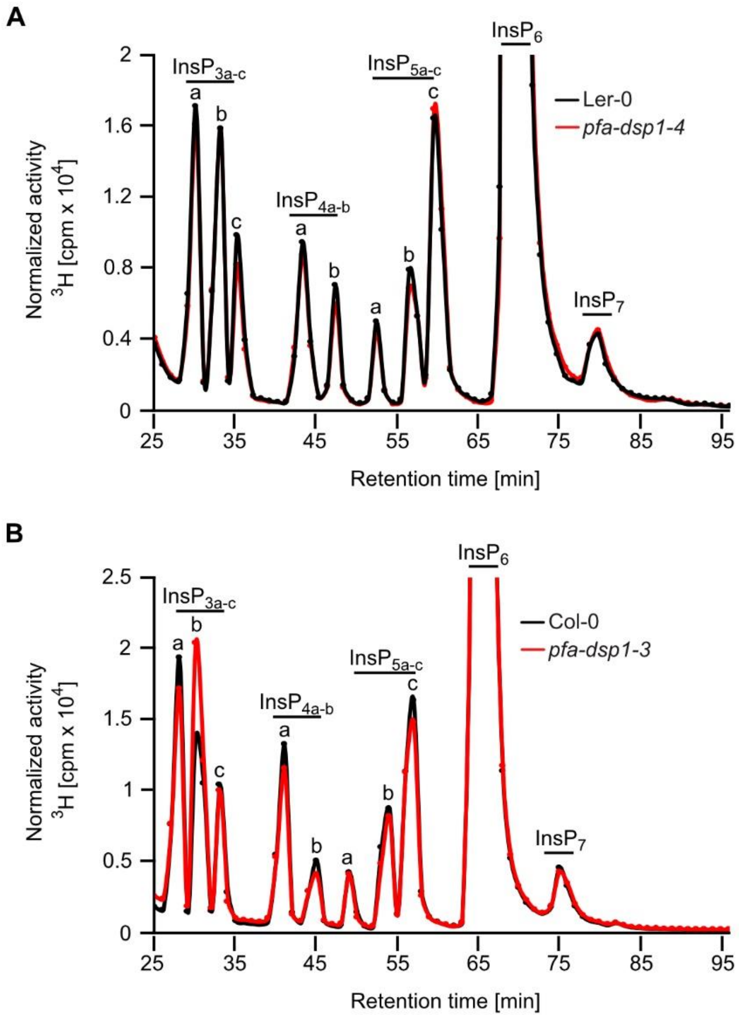


**Figure S10: Single mutant *Arabidopsis pfa-dsp1* loss-of-function lines do not display InsP/PP-InsP defects.** Representative SAX-HPLC profiles of 20-days-old wild-type Ler-0 and *pfa-dsp1-4 Arabidopsis* seedlings (A) and of Col-0 and *pfa-dsp1-3 Arabidopsis* seedlings (B) radiolabeled with [$^{3}$H]-*myo*-inositol. All visible peaks are highlighted and assigned to the corresponding InsP species. Based on published chromatographic mobilities [1, 2], $InsP_{4a}$ likely represents Ins(1,4,5,6)$P_4$ or Ins(3,4,5,6)$P_4$, $InsP_{5a}$ likely represents $InsP_5$ [2-OH], $InsP_{5b}$ likely represents $InsP_5$ [4-OH] or its enantiomeric form $InsP_5$ [6-OH], and $InsP_{5c}$ likely represents $InsP_5$ [1-OH] or its enantiomeric form $InsP_5$ [3-OH]. The isomeric natures of $InsP_{3a-c}$, $InsP_{4b}$, $InsP_7$, and $InsP_8$ are unknown.

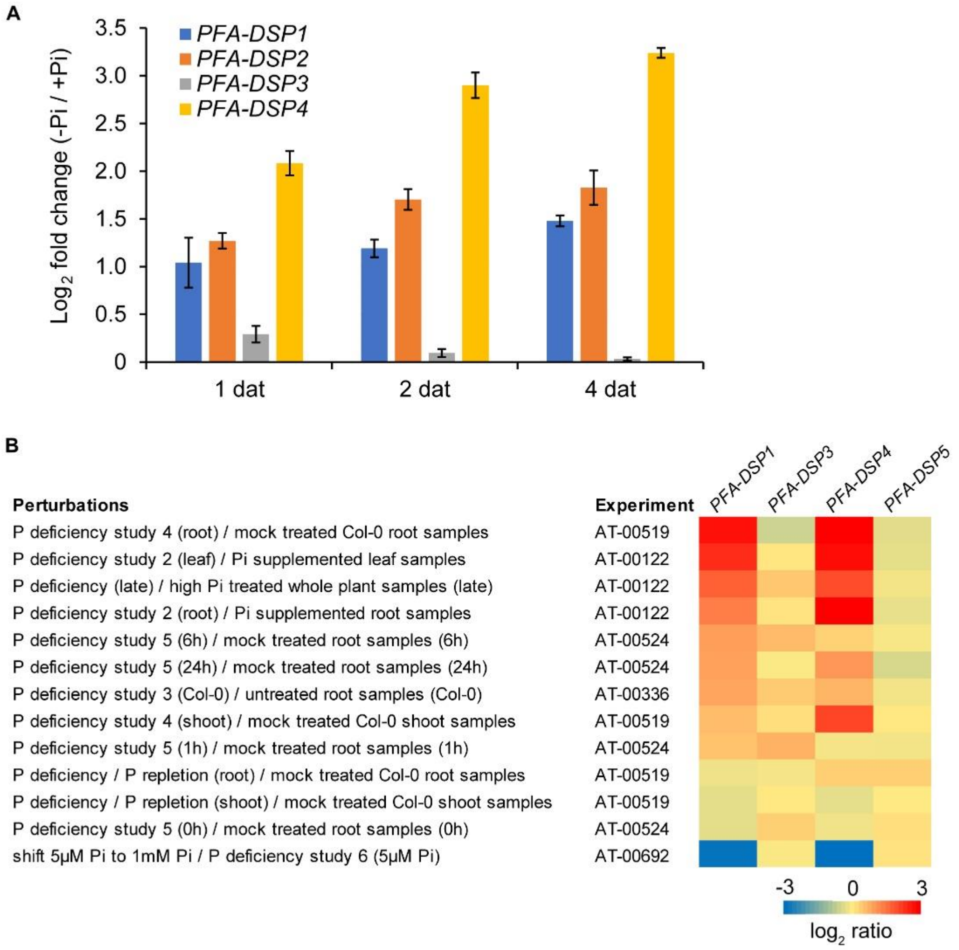


**Figure S11: *Arabidopsis* PFA-DSP1, 2 and 4 are strongly induced by $P_i$ deficiency.** (A) Expression of the indicated *PFA-DSP*s in roots of *Aradidopsis thaliana* (accession Col-0) plants according to a transcriptome experiment with Agilent microarrays [3]; data deposited on e!DAL repository under the accession code https://doi.org/10.5447/IPK/2018/4. No probe for *PFA-DSP5* was present in the microarray chips. Seven-day-old plants pre-cultured on sufficient $P_i$ supply were transferred to fresh solid media containing 625 µM $P_i$ (+Pi) or 100 µM $P_i$ (-Pi). Whole roots were collected at the indicated time points after transfer. Data represent means ± SD (n = 3). (B) Heatmap analysis of *PFA-DSP*s genes in response to the indicated $P_i$ treatmens. No data are presented for *PFA-DSP2* as no probe for this gene is present in Affimetrix chips. Transcriptional data were retrieved and analyzed with Genevestigator (http://www.genevestigator.ethz.ch).

**Table S1: Oligonucleotide sequences.**

| Primer name | Sequence |
|---|---|
| attB1 adapter | GGGGACAAGTTTGTACAAAAAAGCAGGCTTC |
| attB2 adapter | GGGGACCACTTTGTACAAGAAAGCTGGGTC |
| attB2+V5 adapter | GGGGACCACTTTGTACAAGAAAGCTGGGTCTTACGTAGAATCGAGACCGAGGAGAGGG TTAGGGATAGGCTTACCTCCTCCAGATCC |
| attB1_ScSIW14 | AAAAAGCAGGCTTCATGGGTTTATATCAAGCAAAG |
| attB2_ScSIW14s | AAAAAGCAGGCTTCATGGGTTTATATCAAGCAAAG |
| attB2_ScSIW14V5 | CTTACCTCCTCCAGATCCCCATTGTAGAGGCAACCAG |
| attB1_AtPFA-DSP1 | AAAAAGCAGGCTTCATGAAGCTTGTGGAGAAGAC |
| attB2_AtPFA-DSP1ns | AGAAAGCTGGGTCCCTGATGGAACAAGAGAATG |
| attB2_AtPFA-DSP1s | AGAAAGCTGGGTCTTACCTGATGGAACAAGAG |
| attB2_AtPFA-DSP1V5 | ACCTCCTCCAGATCCCCTGATGGAACAAGAGAATG |
| attB1_AtPFA-DSP2 | AAAAAGCAGGCTTCATGAAACTGATTGAGAAGACG |
| attB2_AtPFA-DSP2s | AGAAAGCTGGGTCTTACCTATTGGAGCAAGAAAAAG |
| attB2_AtPFA-DSP2V5 | ACCTCCTCCAGATCCCCTATTGGAGCAAGAAAAAGAC |
| attB1_AtPFA-DSP3 | AAAAAGCAGGCTTCATGTGTTTGATTATGGAAACGG |
| attB2_AtPFA-DSP3s | AGAAAGCTGGGTCTTAAACTCTAGCAGCCTGCG |
| attB2_AtPFA-DSP3V5 | ACCTCCTCCAGATCCAACTCTAGCAGCCTGCGG |
| attB1_AtPFA-DSP4 | AAAAAGCAGGCTTCATGACGTTAGAGAGTTACGCCG |
| attB2_AtPFA-DSP4s | AGAAAGCTGGGTCTCAGTAATCAATAGTATTAGTATACCTCTTGG |
| attB2_AtPFA-DSP4V5 | ACCTCCTCCAGATCCGTAATCAATAGTATTAGTATACCTCTTGG |
| attB1_AtPFA-DSP5 | AAAAAGCAGGCTTCATGGGCTTAATTGTGGATGATG |
| attB2_AtPFA-DSP5s | AGAAAGCTGGGTCTTATCCTTTGGTGGCTTGAGG |
| attB2_AtPFA-DSP5V5 | ACCTCCTCCAGATCCTCCTTTGGTGGCTTGAGG |
| ScSIW14_C214S_F | TCAACCGATACTGATACATTCTAATAGAGGCAAACATAGAAC |
| ScSIW14_C214S_R | GTTCTATGTTTGCCTCTATTAGAATGTATCAGTATCGGTTGA |
| AtPFA-DSP1_C150S_F | GTTCTGATTCATAGTAAGCGAGGC |
| AtPFA-DSP1_C150S_R | GCCTCGCTTACTATGAATCAGAACA |
| ScSIW14pgt_PstI_F | AGCCTGCAGGATGGAGCTGCTCCTGGCTG |
| ScSIW14pgt_EcoRI_R | GAATTCAATATAAAGCGGGAATTTTTTTTTTC |
| AtPFA-DSP1_267_F | ATACTTGTGCCCGGAGCCCT |
| AtPFA-DSP1_373_R | TCACAAATGGCTCCTTGTTGCCT |
| AtTIP41-like_F | TGGTTGGAAGCAGGAAGGGCT |
| AtTIP41-like_R | TGCTGAGACGGCTTGCTCCTGA |
| AtPP2AA3_F_qPCR | TGGTGCTCAGATGAGGGAGA |
| AtPP2AA3_R_qPCR | TAGCACATCTGGGGCACTTG |
| ScSIW14_pUG_F | CTCTTCTGGATCAATTTTTCTTTTTCATCTAAAGTTTAAAAGGAGCAGCTGAAGCTTCGTA CGC |
| ScSIW14_pUG_R | CATCATTTTCGAAGAGACTAGTTACGTAAAGGTAATCACTGTCTACATAGCATAGGCCAC TAGTGGATCTG |
| WiscDsLox_473B10_LP<br>WiscDsLox_473B10_RP | TTGTTTTGCAAAACTGCAAAG<br>TTGCCTTCAATACCAAACTGG |
| P745_WiscDsLox_F | AACGTCCGCAATGTGTTATTAAGTTGTC |
| GT1415_F | CGACTCTCCTCACCTAAAGATTCA |
| GT1415_R | GTTGCCTTCAATACCAAACTGG |
| DS3-1 | ACCCGACCGGATCGTATCGGT |
| SAIL_116_C12_LP | TTGTTTTGCAAAACTGCAAAG |
| SAIL_116_C12_RP | TTGCCTTCAATACCAAACTGG |
| LB1_SAIL_F | GCCTTTTCAGAAATGGATAAATAGCCTTGCTTCC |

[&]Present address: Department of Biomedicine, University of Basel, 4058 Basel, Switzerland
